# Orphan CpG islands boost the regulatory activity of poised enhancers and dictate the responsiveness of their target genes

**DOI:** 10.1101/2020.08.05.237768

**Authors:** Tomás Pachano, Víctor Sánchez-Gaya, María Mariner-Faulí, Thais Ealo, Helena G. Asenjo, Patricia Respuela, Sara Cruz-Molina, Wilfred F. J. van Ijcken, David Landeira, Álvaro Rada-Iglesias

**Affiliations:** Center for Molecular Medicine Cologne (CMMC), University of Cologne, Robert-Koch-Strasse 21, 50931 Cologne, Germany; Institute of Biomedicine and Biotechnology of Cantabria (IBBTEC), CSIC/Universidad de Cantabria, Albert Einstein 22, 39011 Santander, Spain; Centre for Genomics and Oncological Research (GENYO), Avenue de la Ilustración 114, 18016 Granada, Spain.; Department of Biochemistry and Molecular Biology II, Faculty of Pharmacy, University of Granada, Granada, Spain; Instituto de Investigación Biosanitaria ibs.GRANADA, Hospital Virgen de las Nieves, Granada, Spain; Max Planck Institute for Molecular Biomedicine, Roentgenstrasse 20, 48149 Muenster, Germany.; Erasmus Medical Center, University of Rotterdam, Rotterdam, the Netherlands.; Cologne Excellence Cluster for Cellular Stress Responses in Aging-Associated Diseases (CECAD), University of Cologne, Joseph-Stelzmann-Strasse 26, 50931 Cologne, Germany

## Abstract

CpG islands (CGIs) represent a distinctive and widespread genetic feature of vertebrate genomes, being associated with ∼70% of all annotated gene promoters^1^. CGIs have been proposed to control transcription initiation by conferring nearby promoters with unique chromatin properties^2–4^. In addition, there are thousands of distal or orphan CGIs (oCGIs) whose functional relevance and mechanism of action are barely known^5–7^. Here we show that oCGIs are an essential component of poised enhancers (PEs)^8, 9^ that boost their long-range regulatory activity and dictate the responsiveness of their target genes. Using a CRISPR/Cas9 knock-in strategy in mESC, we introduced PEs with or without oCGIs within topological associating domains (TADs) harbouring genes with different types of promoters. By evaluating the chromatin, topological and regulatory properties of the engineered PEs, we uncover that, rather than increasing their local activation, oCGIs boost the physical and functional communication between PEs and distally located developmental genes. Furthermore, we demonstrate that developmental genes with CpG rich promoters are particularly responsive to PEs and that such responsiveness depends on the presence of oCGIs. Therefore, our work unveils a novel role for CGIs as genetic determinants of the compatibility between genes and enhancers, thus providing major insights into how developmental gene expression programs are deployed under both physiological and pathological conditions^10–12^.

---

The establishment of cell-type specific gene expression programs during vertebrate development is largely dependent on enhancers^13–15^, a group of distal *cis*-regulatory elements containing clusters of transcription factor binding sites (TFBS) that can control gene expression in a distance and orientation independent manner. The regulatory properties of enhancers have been mostly investigated using transgenic reporter assays^16^, in which the enhancer activity of selected sequences is evaluated by measuring their capacity to activate transcription of a reporter gene from a minimal promoter. In these assays, the investigated enhancers are placed at relatively small distances from the reporter genes and using a rather limited set of minimal promoters. On the other hand, the regulatory activity of enhancers can be specifically directed towards their target genes by insulators, which can prevent enhancers from ectopically activating non-target genes^10^. In vertebrates, insulators are typically bound by CTCF, which, together with Cohesin, can form long-range chromatin loops that demarcate the boundaries of regulatory domains (*i.e.* TADs^17^, insulated neighborhoods^18^) and limit enhancer activity towards genes located within the same domain^17, 19^. Altogether, current models of enhancer function implicitly assume that enhancers and genes can effectively communicate with each other, regardless of distance or sequence composition, as far as they are located within the same regulatory domain^10, 11^. However, these models have been challenged by recent studies showing that the loss or structural disruption of regulatory domains does not always lead to changes in gene expression or enhancer-gene communication^20–23^. Therefore, additional factors, besides being in the same regulatory domain, might contribute to the compatibility between genes and enhancers. In this regard, massively parallel reporter assays uncovered that enhancer responsiveness in flies depends on the sequence composition of core promoter elements^24, 25^. However, it is currently unknown whether other genetic factors, such as distance or enhancer sequence composition, can also contribute to such responsiveness. Uncovering the rules controlling the responsiveness of genes to enhancers is essential to understand and predict the pathological consequences of human structural variation^12^.

Focusing on a set of highly conserved developmental enhancers, known as PEs, here we have used a synthetic engineering approach to systematically dissect the genetic rules controlling gene-enhancer compatibility. We previously showed that PEs are essential for the induction of major anterior neural genes upon ESC differentiation^9^. Before becoming active in anterior neural progenitors (AntNPC), PEs are already bookmarked in embryonic stem cells (ESC) with unique chromatin and topological features, including binding by polycomb-group protein complexes (PcG) and pre-formed contacts with their target genes^8, 9^. Compared to other enhancer types, PEs display a distinctive genetic composition that includes not only clusters of TFBS but also nearby CGIs. CGIs are a prevalent feature of gene promoters in vertebrates, where they are believed to provide a permissive chromatin state that facilitates transcription initiation^2–4^. However, only half of the CGIs found in the mouse and human genomes are associated to promoters (pCGIs)^1, 5^; while the other half, typically known as oCGI, remain poorly studied. oCGIs have been proposed to act as alternative gene promoters^6, 7^ or as highly active and conserved enhancers with limited tissue specificity^5, 26, 27^. Nevertheless, these proposed functions are largely based on correlative observations and the mechanisms whereby oCGIs might contribute to gene expression control remain fully unknown. Here we show that oCGIs act as long-range boosters of PE regulatory activity that enable functional communication between PEs and developmental genes. Notably, we also show that such communication is efficiently established with CpG-rich but not with CpG-poor promoters, suggesting that distal sequences, namely oCGIs, can also contribute to enhancer responsiveness and gene expression specificity. Therefore, our work provides major insights into the genetic basis of gene-enhancer compatibility and, thus, into how gene expression programs can be specifically and precisely deployed during development as well as pathologically disrupted by structural variants^10–12^.

## Genetic properties of PE-associated oCGIs

We previously reported that PEs are commonly located in proximity to computationally predicted CGIs^9^. However, computational models underestimate the abundance of CGIs, especially those that are distally located from transcription start sites (TSS)^28^. Therefore, we used biochemically identified non-methylated islands (NMI) to more accurately study the association between PEs and oCGIs^28^ and found that ∼80% of PEs are located within 3Kb of a NMI (Fig. 1a). Next, we genetically compared the NMI associated to PEs (PE-NMI) with those found in proximity of the TSS of developmental genes (devTSS-NMI) (see Methods). In line with the previous characterization of intergenic NMI^5, 28^, PE-NMI are shorter in length and present lower CpG density than devTSS-NMI (Fig. 1b). Similar results were observed using an independent set of biochemically defined CGI (*i.e.* CAP-CGI) (Extended Data Fig. 1)^5^. Thus, PEs are pervasively found in proximity of CGIs genetically distinct from those associated with gene promoters.

**Fig. 1.**
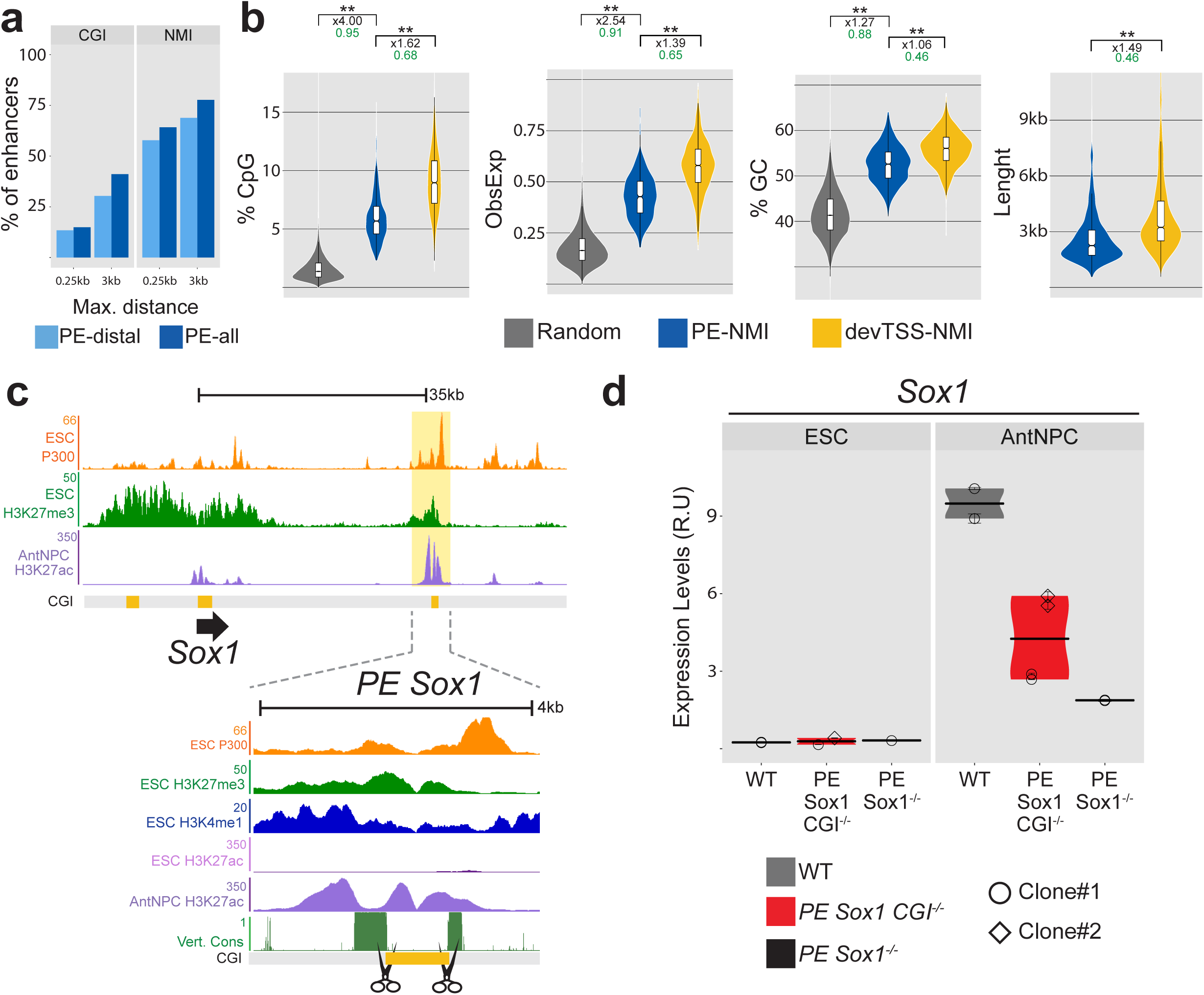
Genetic properties and functional relevance of the orphan CGIs associated with PE. a, Percentage of poised enhancers (PEs) found within the indicated maximum distances (0.25Kb or 3Kb) to a computationally-defined CGI according to the following criteria: GC content > 50%; Length > 200 bp; CpG (left panel) or a NMI identified with the Bio-CAP assay^26^ (right panel). PEs categorized as PE-distal correspond to those located more than 10 Kb away from their closest gene. b, Comparison of the CpG percentage, observed/expected CpG ratio, GC percentage and sequence length between random regions (see Methods), NMIs associated with poised enhancers (PE-NMI; blue) and NMIs associated with the TSS of developmental genes (devTSS-NMI; yellow; see Methods). On top of each plot, the asterisks indicate P-values calculated using unpaired Wilcoxon tests with Bonferroni correction for multiple testing (** = p.val < 1e^-10^; * p.val < 0.05); the numbers in black indicate the median fold changes between the indicated groups; the green numbers indicate non-negligible Cliff Delta effect sizes. c, The upper panel shows ChIP-seq data^9^ from mESC (P300 and H3K27me3) and AntNPC (H3K27ac) at the *Sox1* locus. The *PE Sox1(+35)* is highlighted in yellow. The lower panel shows a close-up view of the *PE Sox1(+35)* with additional epigenomic and genomic data. The represented CGIs correspond to those computationally defined in the UCSC browser according to the following criteria: GC content > 50%; Length > 200 bp; CpG Observed to expected ratio > 0.6. Vert. Cons.= vertebrate PhastCons. d, The expression of *Sox1* was investigated by RT-qPCR in mESCs (left panel) and AntNPC (right panel) that were either WT (grey), homozygous for a deletion of the CGI associated to the *PE Sox1(+35)* (*PE Sox1 CGI^-/-^;* red) or homozygous for the complete *PE Sox1(+35)* deletion^9^ (*PE Sox1^-/-^;* black). Two different *PE Sox1 CGI^-/-^* mESC clones (circles and diamonds) and one *PE Sox1^-/-^* clone were studied. For each mESC clonal line, two technical replicates of the AntNPC differentiation were performed. The plotted expression values of each clone correspond to the average and standard deviation (error bars) from three RT-qPCR technical replicates. Expression values were normalized to two housekeeping genes (*Eef1a* and *Hprt*). The results of an independent biological replicate for this experiment are shown in Extended Data Fig. 2c.

## oCGIs are necessary for the regulatory function of PEs

To start evaluating the regulatory role of oCGIs in the context of PEs, we first generated mESC lines with a homozygous deletion of the oCGI associated with a previously characterized PE (i.e. *PE Sox1(+35)*^9^) that controls the expression of *Sox1* in neural progenitors (*PE Sox1(+35)-CGI^-/-^* mESC; Fig. 1c and Extended Data Fig. 2a). ChIP-qPCR experiments in mESC showed that the deletion of the oCGI severely reduced H3K27me3 levels around the *PE Sox1(+35)* region (Extended Data Fig. 2b), thus demonstrating that oCGIs are necessary for the recruitment of PcG to PE. Next, to investigate how the loss of the oCGI might affect *PE Sox1(+35)* function, we measured *Sox1* expression in WT, *PE Sox1(+35)-CGI^-/-^* and *PE Sox1(+35)^-/-^* mESC as well as upon their differentiation into AntNPC. In mESC, neither the deletion of the oCGI nor of the whole *PE Sox1(+35)* affected *Sox1* expression (Fig. 1d; Extended Data Fig. 2c). Together with our previous work^9^, these results further support that PEs do not act as silencers in ESC and that PcG recruitment is not required to keep PEs is an inactive sate. However, upon differentiation into AntNPC, the deletion of the oCGI reduced *Sox1* induction by >2-fold (compared to >4-fold reduction in cells with the full PE deletion) (Fig. 1d; Extended Data Fig. 2c). These findings suggest that, rather than protecting PEs from a premature activation through the recruitment of PcG, oCGIs might positively influence the *cis*-activation capacity of PEs.

## Dissection of the regulatory logic of PEs using a genetic engineering approach

Loss-of-function approaches based on genetic deletions have several limitations in order to conclusively assess the function of oCGIs within PEs: (i) oCGIs can be difficult to delete individually, as they frequently overlap with the nearby TFBS (Extended Data Fig. 3a); (ii) the deletion of the oCGI can affect PE activity by altering the distance between TFBS^29^; (iii) PE target genes typically display complex regulatory landscapes in which multiple enhancers might control gene expression^30^, thus potentially masking the regulatory function of individual oCGIs; (iv) experiments based on the complete loss of CGI-bound proteins (e.g. PcG, Trithorax-group proteins (TrxG)^31^) can elicit global transcriptional, epigenomic and/or topological changes in mESC and, thus, indirectly alter PE loci.

To systematically dissect the contribution of oCGIs to the regulatory function of PEs, we designed a genetic engineering approach to generate mESC in which the components of selected PEs can be modularly inserted (*i.e.* TFBS, oCGI or TFBS+oCGI) into a fixed genomic location (Fig. 2a). We reasoned that by selecting insertion sites located within TADs containing developmental genes not expressed in ESC or AntNPC and, thus, without active enhancers in these cell types, any changes in the expression of the selected genes could be attributed solely to the inserted PE sequences. Furthermore, by inserting the same PE components within TADs containing developmental genes with different types of promoters (e.g. CpG-rich or CpG-poor), we could interrogate whether gene promoters differ in their responsiveness to PEs. To implement this approach, we used a CRISPR-Cas9 knock-in system^32^ to initially insert the *PE Sox1(+35)* components (*i.e. PE Sox1(+35)TFBS* (465bp), *PE Sox1(+35)CGI* (893bp) or *PE Sox1(+35)TFBS+CGI* (1370bp)) approximately 100 Kb downstream of *Gata6* (Fig. 2a), a gene with multiple CGIs around its promoter region and lowly expressed in both ESC and AntNPC (0.4 and 0.06 FPKMs, respectively^9^). The selected insertion site was not conserved and did not overlap or was close to any CTCF binding site, thus minimizing the risk of disrupting a regulatory element. Using this strategy, we successfully established two homozygous mESCs clones for each of the *PE Sox1(+35)* insertions described above (Extended Data Fig. 3b). Next, we measured *Gata6* expression in the previous mESC lines as well as upon their differentiation into AntNPC. In undifferentiated ESC, we found that none of the engineered *PE Sox1(+35)* combinations affected *Gata6* expression (Fig. 2b and Extended Data Fig. 3c). Strikingly, upon differentiation into AntNPC, *Gata6* was strongly induced in cells with the *PE Sox1(+35)TFBS+CGI* insertion (∼50-fold *vs* WT). In contrast, cells with the *PE Sox1(+35)TFBS* insert displayed considerably milder *Gata6* induction (∼7-fold *vs* WT), while the insertion of the *PE Sox1(+35)CGI* had not effect on *Gata6* expression (Fig. 2b and Extended Data Fig. 3c). Therefore, these initial results suggest that, although the oCGI do not have *cis*-regulatory activity on their own, they can dramatically boost the regulatory activity of their nearby TFBS.

**Fig. 2.**
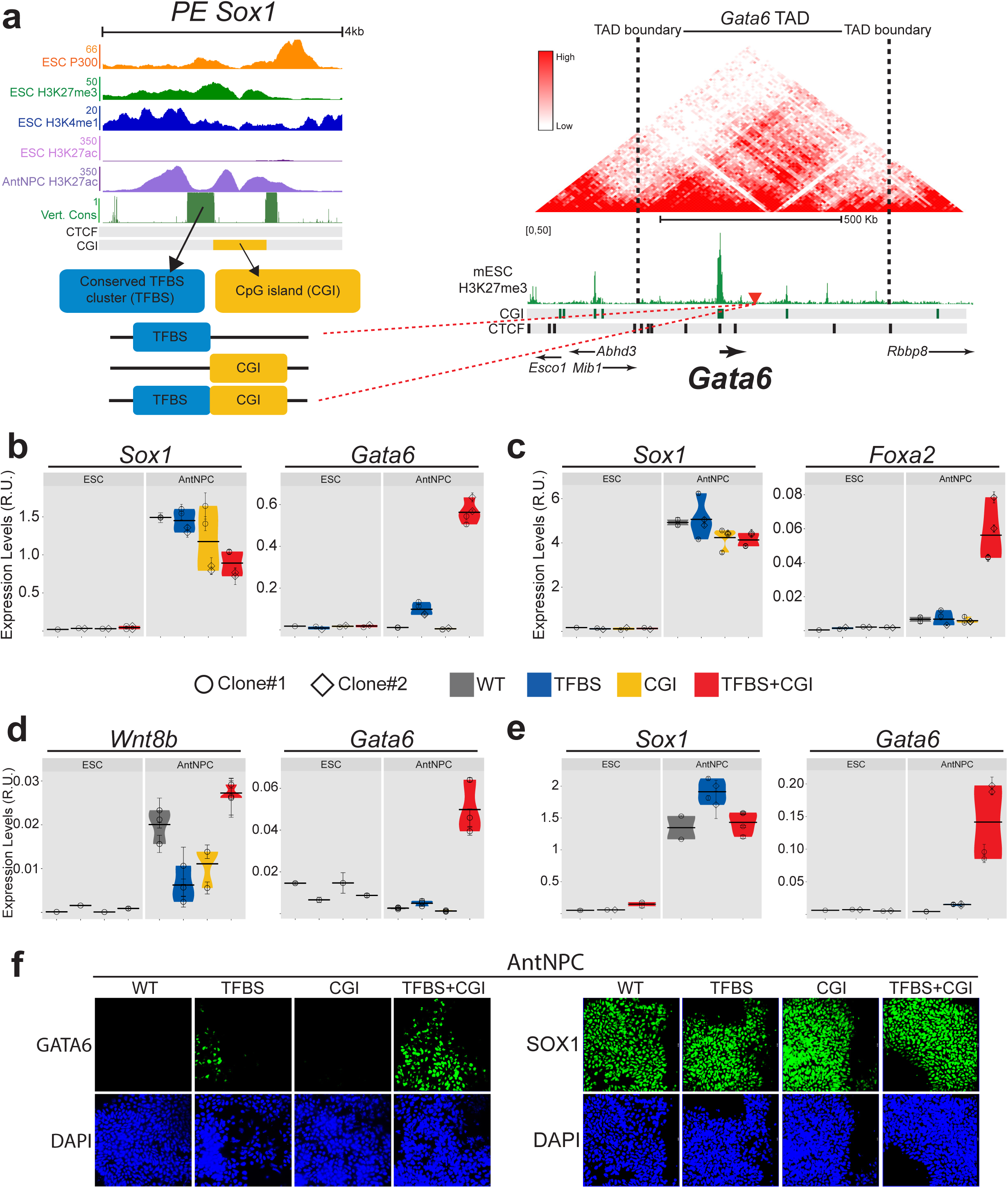
Modular engineering of PEs reveals major regulatory functions for orphan CGIs. a, Strategy used to insert the *PE Sox1(+35)* components into the *Gata6-*TAD. The upper left panel shows a close-up view of the epigenomic and genetic features of the *PE Sox1(+35)* (Vert. Cons.= vertebrate PhastCons). The represented CGI was computationally defined according to the following criteria: GC content > 50%; Length > 200 bp; CpG Observed to expected ratio > 0.6. The lower left panel shows the three combinations of *PE Sox1(+35)* modules (i.e. (i) *PE Sox1(+35)TFBS; (ii) PE Sox1(+35)CGI; (iii) PE Sox1(+35)TFBS&CGI)* inserted into the *Gata6* TAD. The right panel shows the TAD in which *Gata6* is included (i.e. *Gata6-*TAD) according to publically available Hi-C data^33, 34^; TAD boundaries are denoted with dotted lines; H3K27me3 ChIP-seq signals in mESC are shown in green^9^; CTCF binding sites in ESC^35^ are shown as black rectangles; CGIs are indicated as green rectangles; the red triangle indicates the integration site of the *PE Sox1(+35)* modules approximately 100 Kb downstream of *Gata6*. b-e, The expression of *Gata6 (b,d and e), Foxa2 (c), Sox1 (b, c and e) and Wnt8b (c)* was measured by RT-qPCR in mESCs (left panels) and AntNPC (right panels) that were either WT (grey) or homozygous for the insertions of the different *PE Sox1(+35)* (b and c) or *PE Wnt8b(+21)* (d) modules (i.e. TFBS (blue), CGI (yellow), TFBS+CGI (red)). In (e), *PE Sox1(+35)TFBS* was inserted alone (blue) or in combination with an artificial CGI (red) into the *Gata6-*TAD. For the cells with the PE module insertions, two different clonal cell lines (circles and diamonds) were studied in each case. For each cell line, two technical replicates of the AntNPC differentiation were performed. The plotted expression values for each clone correspond to the average and standard deviation (error bars) from three RT-qPCR technical replicates. Expression values were normalized to two housekeeping genes (*Eef1a* and *Hprt*). The results of an independent biological replicate for each experiment are shown in Extended Data Fig. 3-5. f, The expression patterns of the GATA6 (left panel) and SOX1 (right panel) proteins were investigated by immunofluorescence in AntNPCs that were either WT or homozygous for the insertion of the different *PE Sox1(+35)* modules (i.e. TFBS, CGI, TFBS+CGI) in the *Gata6-*TAD. Nuclei were stained with DAPI.

**Fig. 3.**
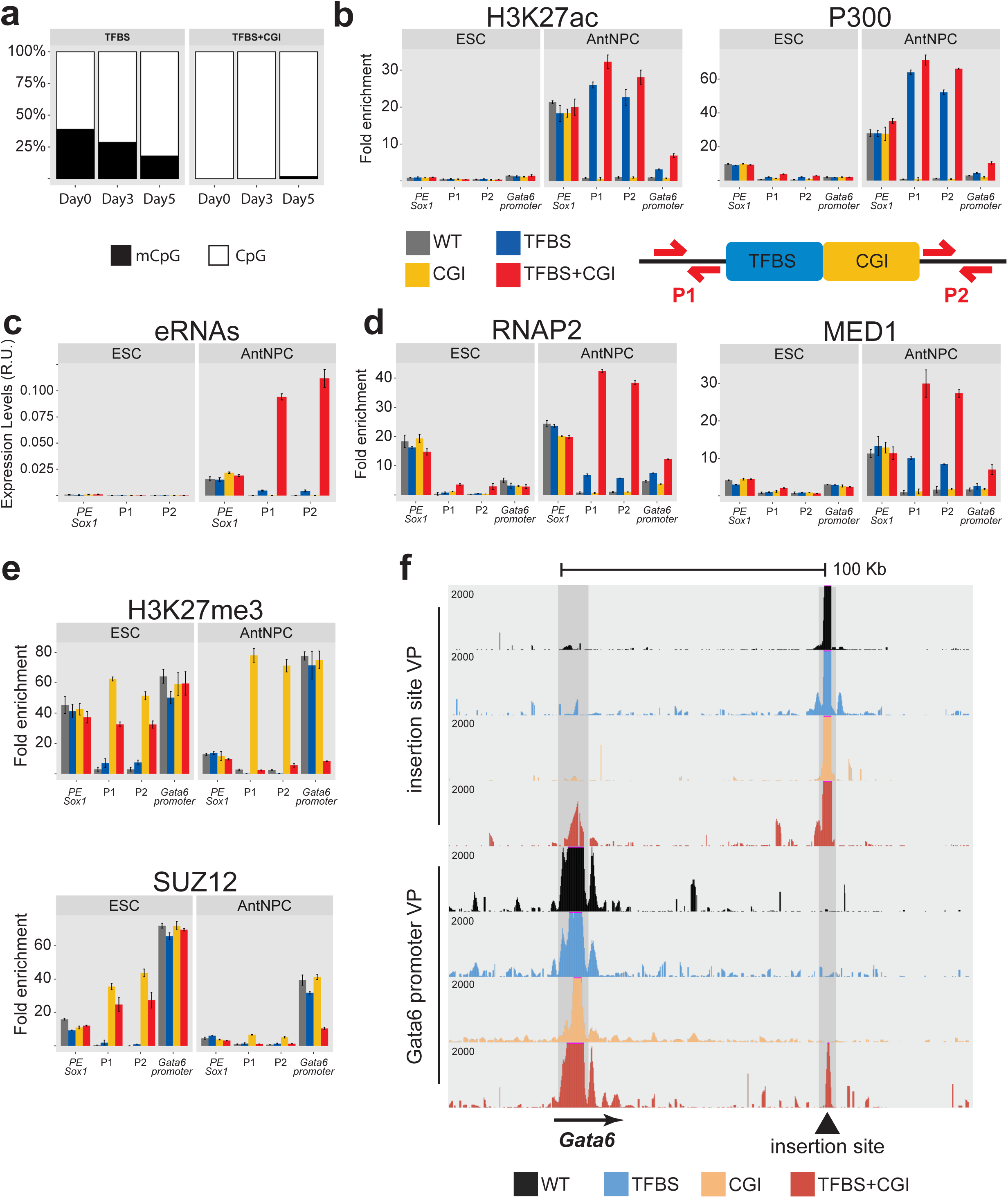
Characterization of the epigenetic, topological and regulatory features of the *PE Sox1(+35)* modules engineered within the Gata6-TAD. a, Bisulfite sequencing analyses during mESC (Day0) to AntNPC (Day5) differentiation of cell lines with the *PE Sox1(+35)TFBS* or *PE Sox1(+35)TFBS+CGI* modules inserted in the *Gata6-*TAD. DNA methylation levels were measured using a forward bisulfite primer upstream of the insertion site and a reverse primer inside the TFBS module (see Methods). **b**, H3K27ac and P300 levels at the endogenous *PE Sox1(+35)*, the *Gata6-*TAD insertion site (primer pairs P1 and P2) and the *Gata6* promoter were measured by ChIP-qPCR in mESCs (left panels) and AntNPC (right panels) that were either WT (gray) or homozygous for the insertions of the different *PE Sox1(+35)* modules (i.e. TFBS (blue), CGI (yellow), TFBS+CGI (red)). ChIP-qPCR signals were normalized against two negative control regions (Supplementary Data 1). Error bars correspond to standard deviations from technical triplicates. The location of the primers P1 and P2 around the *Gata6* TAD insertion site is represented as red arrows in the lower diagram. **c**, eRNA levels at the endogenous *PE Sox1(+35)* and the *Gata6-*TAD insertion site (primers P1 and P2) were measured by RT-qPCR in mESCs (left panels) and AntNPC (right panels) that were either WT (gray) or homozygous for the insertions of the different *PE Sox1(+35)* modules (i.e. TFBS (blue), CGI (yellow), TFBS+CGI (red)). The plotted expression values correspond to the average and standard deviation (error bars) from three RT-qPCR technical replicates. Expression values were normalized to two housekeeping genes (*Eef1a* and *Hprt*). **d,e**, RNAP2 and MED1 (d) or H3K27me3 and SUZ12 (e) levels were measured by ChIP-qPCR as described in (b). **f**, 4C-seq experiments were performed using the *Gata*6-TAD insertion site (upper panels) or the *Gata6* promoter (lower panels) as viewpoints in mESCs that were either WT (black) or homozygous for the insertions of the different *PE Sox1(+35)* modules (i.e. TFBS (blue), CGI (yellow), TFBS+CGI (red)).

## oCGIs boost the regulatory activity of PEs

To evaluate whether the previous observations could be generalized, we used the same genetic engineering approach to generate two additional group of transgenic ESC lines: (i) the same *PE Sox1(+35)* components were inserted within the *Foxa2*-TAD (∼100 Kb downstream of the *Foxa2* TSS, which also contains several CGIs and is inactive in ESC and AntNPC); (ii) the *PE Wnt8b(+21)*^9^ components (*i.e. PE Wnt8b(+21)TFBS*, *PE Wnt8b(+21)CGI* and *PE Wnt8b(+21)TFBS+CGI*) were inserted within the *Gata6-*TAD (∼100 Kb downstream of the *Gata6*-TSS) (Extended Data Fig. 4a-d). Importantly, in both groups of cell lines we again observed that the TFBS+CGI combinations were able to strongly boost the TFBS *cis*-regulatory activity in AntNPC (13-and 19-fold in comparison with WT for *Foxa2* and *Gata6*, respectively; Fig. 2c,d and Extended Data Fig. 4e,f); while the *TFBS* or the oCGI alone lead to either no or minor gene inductions (Fig. 2c,d and Extended Data Fig. 4e,f). Next, we also wanted to investigate whether the boosting capacity of the oCGI could be attributed to other type of regulatory information beyond their CpG-richness (*e.g.* binding sites for TFs). For this purpose, we designed an artificial CGI (aCGI^36^; see Methods) and inserted it together with the *PE Sox1(+35)TFBS* at the *Gata6*-TAD (*PE Sox1(+35)TFBS+aCGI;* Extended Data Fig. 5a,b). Notably, RT-qPCR experiments in AntNPC showed that *Sox1(+35)TFBS+aCGI* cells expressed considerably higher levels of *Gata6* than *Sox1(+35)TFBS* cells (Fig. 2e and Extended Data Fig. 5c), suggesting that the CpG-richness of oCGIs is sufficient to boost the regulatory activity of PEs.

The boosting properties of oCGIs might be attributed to a premature induction of the target gene, an increase in the number of cells in which the target gene becomes induced and/or an increase in the expression levels within individual cells. To address this, we focused on those

cell lines containing the different *PE Sox1(+35)* components inserted within the *Gata6-*TAD. Upon differentiation of the *Sox1(+35)TFBS+CGI* ESC into AntNPC, *Gata6* did not become significantly induced until Day4, thus perfectly matching the expression dynamics of *Sox1* (the endogenous target of *PE Sox1(+35)*) and arguing against premature target gene induction due to the presence of the oCGI (Extended Data Fig. 6). Next, we performed immunofluorescence assays to visualize GATA6 and SOX1 protein levels in both mESC and AntNPC. In agreement with its widespread expression in neural progenitors^37^, SOX1 became strongly and homogenously induced in AntNPC derived from all the evaluated cell lines (Fig. 2f). Notably, GATA6 was also induced in most of the AntNPC derived from PE *Sox1(+35)TFBS+CGI* mESC (Fig. 2f), thus closely recapitulating the SOX1 expression pattern. In contrast, the *PE Sox1(+35)TFBS* insert resulted in a considerably noisier and more heterogeneous expression of GATA6, while no GATA6 could be detected in *PE Sox1(+35)CGI* cells. These results suggest that, during pluripotent cell differentiation, oCGIs increase the number of cells in which the PE target genes get induced, thus potentially leading to increased gene expression precision^38^.

Taken together, we conclude that oCGIs are an essential component of PEs that might endow them with privileged regulatory properties (e.g. gene expression precision). Notably, our data also suggest that developmental genes with pCGI (e.g. *Gata6* and *Foxa2*) are intrinsically responsive to PEs.

## The boosting capacity of oCGIs does not involve the local activation of PEs

Having identified oCGIs as boosters of PE regulatory activity, we then investigated the mechanisms whereby the oCGI could exert such function, focusing on the mESC lines in which the *PE Sox1(+35)* components were inserted within the *Gata6*-TAD. CGIs are typically devoid of CpG methylation and display low nucleosomal density, which might provide a chromatin environment permissive for TF binding and transcription initiation^39–43^. However, these epigenetic features have been mostly investigated in the context of pCGI. To investigate whether oCGIs can similarly influence the chromatin environment of PEs, we first performed bisulfite sequencing experiments in *PE Sox1(+35)TFBS+CGI* and *PE Sox1(+35)TFBS* cells to measure CpG methylation within the TFBS module. Remarkably, in undifferentiated mESC, the TFBS sequences acquired intermediate CpG methylation levels when inserted alone, while becoming completely unmethylated when combined with the oCGI (Fig. 3a and Extended Data Fig. 7a). Next, we performed FAIRE assays to investigate whether the oCGI could similarly impact chromatin accessibility within the engineered PEs (Extended Data Fig. 7b). In contrast to the dramatic effects observed for CpG methylation, the oCGI only moderately increased chromatin accessibility whether inserted alone or in conjunction with the TFBS. To simultaneously measure nucleosome occupancy and CpG methylation levels at the inserted TFBS with single-DNA molecule resolution^44^, we also performed NOME-PCR assays in *PE Sox1(+35)TFBS+CGI* and *PE Sox1(+35)TFBS* mESC. These experiments confirmed that oCGIs protect nearby TFBS from CpG methylation without having a major effect on chromatin accessibility (Extended Data Fig. 7c,d). Furthermore, upon differentiation into AntNPC, the TFBS also got progressively demethylated in the *PE Sox1(+35)TFBS* cells (Fig. 3a and Extended Data Fig. 7a), suggesting that, even in the absence of an oCGI, TFs can access and activate PEs in AntNPC^45^. To test this prediction, we performed ChIP-qPCR experiments to measure p300 binding and H3K27ac levels, two major hallmarks of active enhancers^46, 47^, around the inserted *PE Sox1(+35)* constructs. Interestingly, these experiments showed that in AntNPC, the PEs containing the TFBS alone or together with the oCGI became strongly and similarly enriched in H3K27ac and P300 (Fig. 3b). Overall, these results indicate that the boosting capacity of the oCGI cannot be simply attributed to their local effects on the chromatin properties of PEs. Nevertheless, the CpG hypomethylation that the oCGI confer to PEs might be related to non-regulatory functions, such as preventing C-to-T mutations ^48^.

## oCGIs increase the physical and functional communication between PEs and their target genes

Another distinctive hallmark of active enhancers is the production of short bidirectional transcripts termed enhancers RNAs (eRNAs), which might be a better predictor of enhancer activity than H3K27ac^49–52^. Remarkably, RT-qPCR analyses in AntNPC showed a >20-fold increase in eRNA levels around the *Sox1(+35)TFBS+CGI* insert in comparison with the *PE Sox1(+35)TFBS* alone (Fig. 3c), thus in full agreement with the effects that these two inserts have on *Gata6* expression. Moreover, additional ChIP-qPCR experiments revealed that, upon AntNPC differentiation, the *Sox1(+35)TFBS+CGI* insert became highly enriched in RNA Polymerase II (RNAP2) and Mediator, a protein complex that acts as a functional bridge transmitting regulatory information from enhancers to promoters^53^(Fig. 3d). In contrast, the binding of these proteins to the *Sox1(+35)TFBS* and *Sox1(+35)CGI* inserts was either considerably weaker or undetectable, respectively (Fig. 3d). Similarly, the recruitment of RNA Pol2 and Mediator at the *Gata6* promoter was also stronger in AntNPC derived from the *PE Sox1(+35)TFBS+CGI* mESC in comparison with the other cell lines (Fig. 3d). All together, these results suggest that oCGIs increase the functional communication between PEs and their target genes, resulting in elevated levels of both eRNAs and mRNAs.

In their inactive state, PEs are enriched in histone modifications (i.e. H3K27me3 and H3K4me1) and are bound by protein complexes (e.g. PcG) that have been previously implicated in the establishment of long-range chromatin interactions^8, 9, 54–59^. Therefore, we wondered if oCGIs could be implicated in the establishment of the PEs unique chromatin signature and thereby facilitate the physical communication between PEs and their target genes. To investigate this possibility, we first performed ChIPs for H3K4me1, H3K4me3 and H3K27me3 in the mESC lines containing the different *PE Sox1(+35)* components within the *Gata6*-TAD (Fig. 3e and Extended Data Fig. 8a,b). H3K4me1 was weakly and similarly enriched around the *PE Sox1(+35)* inserts containing the TFBS with or without the oCGI, while no enrichment was observed for the oCGI insert alone (Extended Data Fig. 8a). Therefore, H3K4me1 deposition seems to be dependent on the TFBS rather than on the oCGI. On the other hand, H3K4me3 was not enriched in any of the evaluated mESC lines (Extended Data Fig. 8a), indicating that oCGIs, perhaps due to their distinct genetic features (Fig. 1), do not adopt the same chromatin state as pCGI^36^. Most interestingly, H3K27me3 was strongly enriched around the *PE Sox1(+35)* inserts containing the oCGI (Fig 3e). ChIPs for additional PcG subunits (i.e. SUZ12, CBX7 and RING1B) and associated histone modifications (i.e. H2AK119ub) further confirmed that oCGIs are sufficient for the recruitment of PcG to PEs (Fig 3e and Extended Fig 8b). Intriguingly, PRC1 recruitment (i.e. H2AK119ub, CBX7 and RING1b) was considerably stronger for the TFBS+oCGI insert than for the insert containing the oCGI alone (Extended Data Fig. 8b).

Since PcG can mediate the establishment of long-range homotypic interactions between distal PcG-bound loci^54–58^, we then investigated the 3D organization of the *Gata6* locus in our engineered mESC lines. 4C-seq experiments using either the *Gata6* promoter or the *PE Sox1(+35)* insertion site as viewpoints revealed that strong PE-*Gata6* contacts were only established in the *PE Sox1(+35)TFBS+CGI* cells (Fig. 3f). The lack of PE-gene contacts in *Sox1(+35)CGI* ESC can be attributed to the weaker recruitment of PRC1 to the PE in these cells (Extended Data Fig. 8b), since long-range interactions between PcG-bound loci are considered to be mediated by PRC1^60–63^. Furthermore, the strong interactions between the *PE Sox1(+35)TFBS+CGI* insert and the *Gata6* promoter were also observed upon differentiation into AntNPC (Extended Data Fig. 8c). In AntNPC, the TFBS+CGI insert lost H3K27me3 and PRC2, thus mirroring the concomitant and strong H3K27ac gains observed in those same cells (Fig. 3e). Therefore, different factors might contribute to the physical communication between PEs and their target genes promoters before and after PE activation.

Overall, our data strongly suggest that, rather than locally facilitating PE activation (*i.e.* p300 recruitment and H3K27ac deposition), oCGIs boost the *cis*-regulatory activity of PEs by bringing genes and enhancers into close spatial proximity and, thus, increasing their functional communication.

## Genes with CpG-poor promoters do not show long-range responsiveness to PEs

All the developmental genes used as PE targets in our genetic engineering approach (Fig. 2) have CpG-rich promoters and are bound by PcG in ESC. Therefore, the high responsiveness of these genes to the PEs could depend not only on the presence of an oCGI within the PE (Fig. 2), but also on CGIs located at the target gene promoters. These CGIs could impose homotypic chromatin states to both PEs and promoters, thus facilitating their long-range physical and functional communication^64^. To test this hypothesis, we inserted the *PE Sox1(+35)* components (*i.e. PE Sox1(+35)TFBS*, *PE Sox1(+35)CGI* or *PE Sox1(+35)TFBS+CGI*) into the *Gria1-TAD,* approximately 100 Kb upstream of the *Gria1* TSS (Fig 4a and Extended Data Fig. 9a). Similarly to *Gata6* and *Foxa2*, *Gria1* is not expressed in either mESC or AntNPC (0 and 0.04 FPKMs, respectively^9^). However, and in contrast to these genes, the *Gria1* promoter does not contain CGIs and is not bound by PcG but fully DNA methylated instead. Remarkably, RT-qPCR analysis of these *Gria1-TAD* cell lines showed that, upon AntNPC differentiation, none of the *PE Sox1(+35)* inserts was able to induce *Gria1* expression (Fig. 4b and Extended Data Fig 9b).

**Fig. 4.**
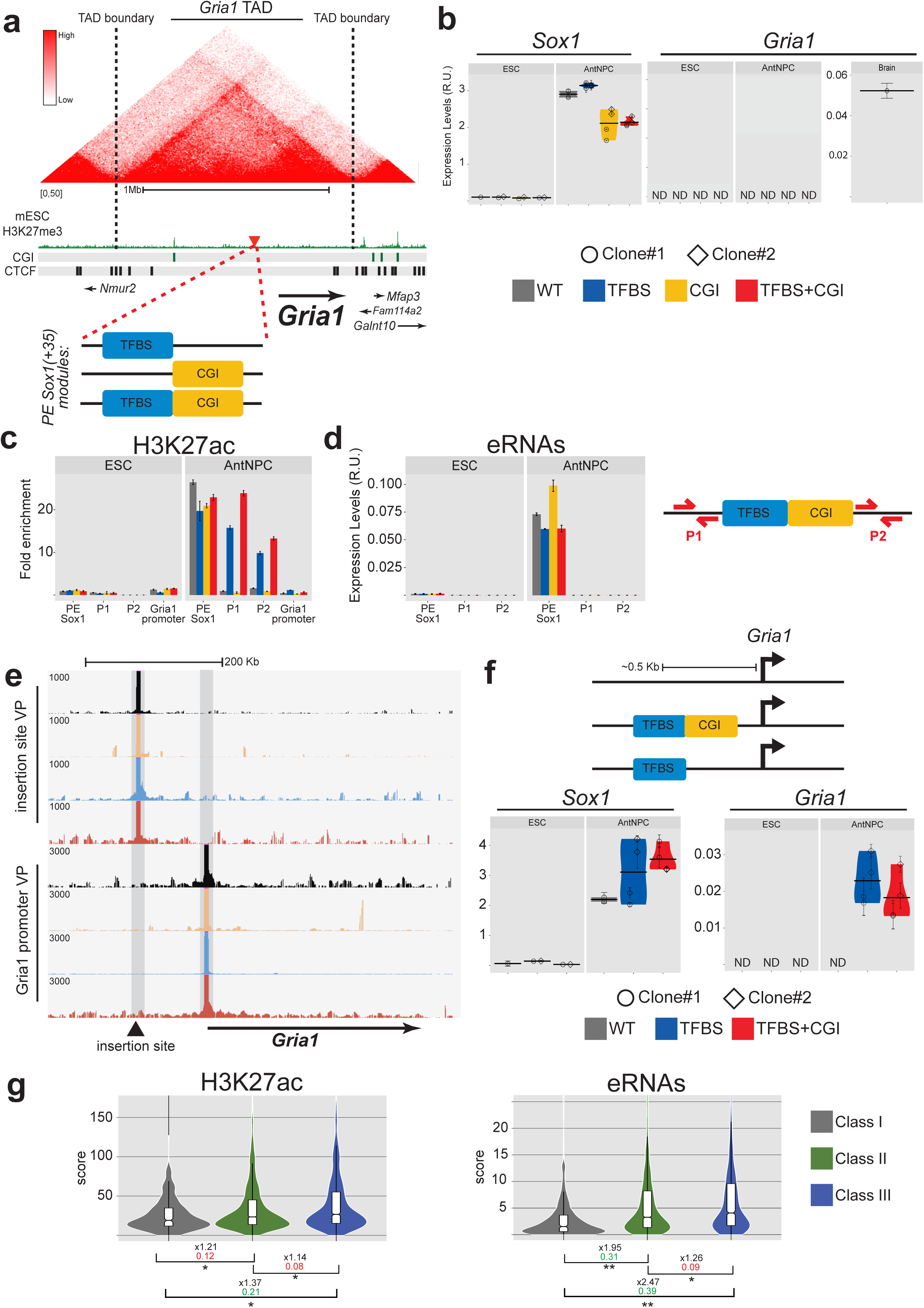
Orphan CGI do not exert their long-range boosting effects over genes with CpG-poor promoters. **a**, Strategy used to insert the *PE Sox1(+35)* components into the *Gria1*-TAD. The lower panel shows the three different combinations of PE modules (i.e. (i) *PE Sox1(+35)TFBS*; (ii) *PE Sox1(+35)CGI*; (iii) *PE Sox1(+35)TFBS&CGI*) inserted into the *Gria1*-TAD. The upper panel shows the TAD in which *Gria1* is included (i.e. *Gria1*-TAD) according to publically available Hi-C data^33, 65^; TAD boundaries are denoted with dotted lines; H3K27me3 ChIP-seq signals in mESC are shown in green^9^; CTCF binding sites in ESC^35^ are shown as black rectangles; CGIs are indicated as green rectangles; the red triangle indicates the integration site of the *PE Sox1(+35)* modules, approximately 100 Kb upstream of *Gria1*. **b**, The expression of *Gria1* and *Sox1* was measured by RT-qPCR in mESCs (left panels) and AntNPC (right panels) that were either WT (grey) or homozygous for the insertions of the different *PE Sox1(+35)* modules (TFBS (blue), CGI (yellow), TFBS+CGI (red)). *Gria1* expression was also measured in mouse embryonic brain to illustrate the quality of the RT-qPCR primers. For the cells with the PE module insertions, two different clonal lines (circles and diamonds) were studied in each case. For each cell line, two technical replicates of the AntNPC differentiation were performed. The plotted expression values for each clone correspond to the average and standard deviation (error bars) from three RT-qPCR technical replicates. Expression values were normalized to two housekeeping genes (*Eef1a* and *Hprt*). The results of an independent biological replicate for each experiment are shown in Extended Data Fig. 9b. c, H3K27ac levels at the endogenous *PE Sox1(+35)*, the *Gria1* TAD insertion site (primer pairs P1 and P2) and the *Gria1* promoter were measured by ChIP-qPCR in mESCs (left panels) and AntNPC (right panels) that were either WT (grey) or homozygous for the insertions of the different *PE Sox1(+35)* modules (i.e. TFBS (blue), CGI (yellow), TFBS+CGI (red)). ChIP-qPCR signals were normalized against two negative control regions (Supplementary Data 1). Error bars correspond to standard deviations from technical triplicates. **d**, eRNAs levels at the endogenous *PE Sox1(+35)* and the *Gria1*-TAD insertion site (primer pair P1 and P2) were measured by RT-qPCR in mESCs (left panels) and AntNPC (right panels) that were either WT (grey) or homozygous for the insertions of the different *Sox1(+35)* modules (i.e. TFBS (blue), CGI (yellow), TFBS+CGI (red)). The plotted expression values correspond to the average and standard deviation (error bars) from three RT-qPCR technical replicates. Expression values were normalized to two housekeeping genes (*Eef1a* and *Hprt*). **e**, 4C-seq experiments were performed using the *Gria1*-TAD insertion site (upper panels) or the *Gria1* promoter (lower panels) as viewpoints in mESCs (left panels) and AntNPC (right panels) that were either WT (black) or homozygous for the insertions of the different *PE Sox1(+35)* modules (i.e. TFBS (blue), CGI (yellow), TFBS+CGI (red)). **f**, The expression of *Gria1* and *Sox1* was measured by RT-qPCR in mESCs (left panels) and AntNPC (right panels) that were either WT or homozygous for the indicated *PE Sox1(+35)* modules (TFBS (blue), TFBS+CGI (red)), which were inserted immediately upstream of the *Gria1* TSS. For the cells with the PE module insertions, two different clonal lines (circles and diamonds) were studied in each case. For each cell line, two technical replicates of the AntNPC differentiation were performed. The plotted expression values for each clone correspond to the average and standard deviation (error bars) from three RT-qPCR technical replicates. Expression values were normalized to two housekeeping genes (*Eef1a* and *Hprt*). The results of an independent biological replicate for each experiment are shown in Extended Data Fig. 10c. **g**, Active enhancers identified in mESC based on the presence of distal H3K27ac peaks (see Methods) were classified into three categories: *Class I* correspond to active enhancers located in TADs containing only poorly expressed genes (<0.5 FPKM); *Class II* correspond to active enhancers located in a TAD with at least one gene with expression levels of >10 FPKM; *Class III* correspond to active enhancers whose closest genes in their same TAD have expression levels of >10 FPKM. The violin plots show the H3K27ac (left) and eRNA (right) levels for each of these active enhancer categories in mESC. On the bottom of each plot, the asterisks indicate P-values calculated using unpaired Wilcoxon tests with Bonferroni correction for multiple testing (** = p.val < 1e^-10^; * p.val < 0.05); the numbers in black indicate the median fold-changes between the indicated groups; the coloured numbers correspond to Cliff Delta effect sizes: negligible (red) and non-negligible (green). The H3K27ac ChIP-seq data was obtained from^9^ and the PRO-seq data to measured eRNA levels was obtained from^66^.

To gain further mechanistic insights into the lack of responsiveness of *Gria1* to the *PE Sox1(+35)*, we then measured DNA methylation, H3K27ac, P300, RNAP2, MED1 and eRNA levels around the inserted *PE Sox1(+35)* constructs. Similarly to what we observed within the *Gata6-TAD*, the TFBS became demethylated in ESC, albeit partially, when combined with the oCGI (Extended Data Fig. 9c). Nevertheless, upon differentiation into AntNPC, the *Sox1(+35)TFBS+CGI* and *Sox1(+35)TFBS* inserts became strongly and similarly enriched in H3K27ac and P300 (Fig. 4c). Therefore, as within the *Gata6-TAD*, the TFBS module was sufficient for the local activation of the PE. However, and in contrast to what we observed in the *Gata6-TAD*, we did not detect eRNA production by any of the *PE Sox1(+35)* inserts (Fig. 4d). Congruently, the recruitment of RNAP2 and MED1 to the *PE Sox1(+35)* was weak regardless of whether the TFBS were alone or together with the oCGI. (Extended Data Fig. 9d). Furthermore, the previous transcriptional regulators were not recruited to the *Gria1* promoter upon differentiation of any of the *Gria1-TAD* ESC lines, thus in full agreement with the lack of *Gria1* induction observed in those cells (Extended Data Fig. 9d).

Together with our experiments within the *Gata6*-TAD, these results show that developmental genes with CpG-rich promoters are particularly responsive to PEs.

## Uncoupling of H3K27ac and eRNA transcription during enhancer activation

The previous genetic engineering experiments within the *Gata6* and *Gria1*-TAD revealed that H3K27ac and eRNA production can be uncoupled from each other and thus signify different steps during PE activation (Fig. 3b,c and Fig. 4c,d). Namely, the accumulation of H3K27ac occurs as PEs become locally activated, while the production of eRNAs, which is coupled to gene transcription, indicates the functional activation of the PEs. Although our findings are in agreement with the original characterization of eRNAs, in which it was reported that eRNA synthesis requires the presence of a target promoter^49^, several recent studies have proposed that enhancers and promoters represent independent transcriptional units^67, 68^. To assess whether our observations can be generalized, we compared eRNA production between three classes of active enhancers using nascent transcription and epigenomic data generated in ESC^66, 69^: (*I*) enhancers located in TADs containing only poorly expressed genes; (*II*) enhancers located in a TAD with at least one gene with high expression levels; (*III*) enhancers whose closest gene within the same TAD has high expression levels (see Methods). Interestingly, class I enhancers showed ∼2 and 2.5-fold lower eRNA levels than *Class II* and *Class III* enhancers, respectively (Fig. 4g), while H3K27ac levels were similar among the three enhancer groups. The differences in eRNA expression between enhancer classes were maintained after correcting for H3K27ac levels or when using independent transcriptional and epigenomic data sets (Extended Data Fig. 11a-c). Although there were some Class I enhancers for which we observed detectable eRNA levels, our results suggest that enhancer and gene transcription are frequently coupled and mutually dependent on enhancer-promoter interactions^49, 50^.

## oCGIs act as long-range regulatory boosters of PEs

The engineering experiments within the *Gata6-TAD* suggest that the responsiveness to PEs involves the physical proximity between PEs and their target genes, which in ESC is likely to be mediated by PcG present at both PEs and promoter regions^9, 60–62, 70^ (Fig. 3f). ChIP experiments in the *Gria1-*TAD ESC lines revealed that, as observed for the *Gata6-*TAD (Fig. 3e), PcG were recruited to the *PE Sox1(+35)* inserts containing an oCGI (Extended Data Fig. 9e). However, 4C-seq analyses in these cells showed that none of the inserted *PE Sox1(+35)* constructs was able to significantly interact with the *Gria1* promoter (Fig. 4e), which, as stated above, does not contain any endogenous pCGI and is not boud by PcG. These results further suggest that the main regulatory function of oCGIs is to facilitate the establishment of long-range PE-gene contacts. To test this prediction, we generated additional ESC lines in which the *PE Sox1(+35)TFBS* or *PE Sox1(+35)TFBS&CGI* constructs were integrated 380 bp upstream of the *Gria1* TSS (Fig. 4f, Extended Data Fig. 10a,b). Remarkably, RT-qPCR analyses in AntNPC derived from the previous ESC lines revealed that both the *PE Sox1(+35)TFBS+CGI* and *PE Sox1(+35)TFBS* inserts were able to induce *Gria1* and that they did so with similar strength (Fig. 4f and Extended Data Fig. 10c). Therefore, the boosting capacity of oCGIs is lost when PEs are located close to gene promoters, strongly suggesting that oCGIs act as long-range boosters of PE regulatory function.

## The combined effects of CGIs and TAD boundaries enable the specific induction of PE target genes

Our data suggest that, in addition to TAD boundaries, the interactions between PE-associated oCGI and pCGI proximal to developmental genes represent an important regulatory layer that ensures that gene expression programs are specifically implemented during embryogenesis. To test this prediction, we decided to genetically engineer the *Six3/Six2* locus due to the following reasons (Fig. 5a): (i) *Six3* and *Six2* are next to each other, yet they are contained within two neighbouring TADs separated by a conserved TAD boundary^71, 72^; (ii) *Six3* and *Six2* display mutually exclusive expression patterns during embryogenesis (e.g. *Six3* in brain; *Six2* in facial mesenchyme)^71^; (iii) the *Six3*-TAD contains a PE (i.e. *PE Six3(-133)*) that controls the induction of *Six3* in AntNPC without any effects on *Six2*^9^; (iv) in mESC, the *PE Six3(-133*) strongly interacts with *Six3* but not with *Six2*^9^ although both genes contain multiple pCGI. Taking all this information into account, we generated mESC with two different genomic rearrangements: (i) a 36 Kb deletion spanning the *Six3/Six2*-TAD boundary (*del36*) and (ii) a 110 Kb inversion that places *Six3* within the *Six2*-TAD and *vice versa* (*inv110*) (Fig. 5a and Extended Data Fig. 12a,b). Interestingly, upon differentiation into AntNPC, *Six2* expression strongly increased in *del36* and *inv110* cells (∼12- and ∼35-fold compared to WT cells, respectively); while the expression of *Six3* expression was dramatically reduced in *inv110* cells (∼77-fold compared to WT cells) and mildly affected in *del36* cells (∼2.5-fold compared to WT cells) (Fig 5b). Furthermore, and in agreement with the previous gene expression changes, 4C-seq experiments in WT and *del36* ESC showed that the deletion of the *Six3/Six2* boundary resulted in increased interactions between *Six2* and the *PE Six3(-133)* (Extended Data Fig. 12c). These results indicate that the PEs can specifically execute their regulatory functions due to the combined effects of TAD boundaries, which provide insulation, and the long-range homotypic interactions between oCGIs and pCGIs, which confer enhancer responsiveness.

**Fig. 5.**
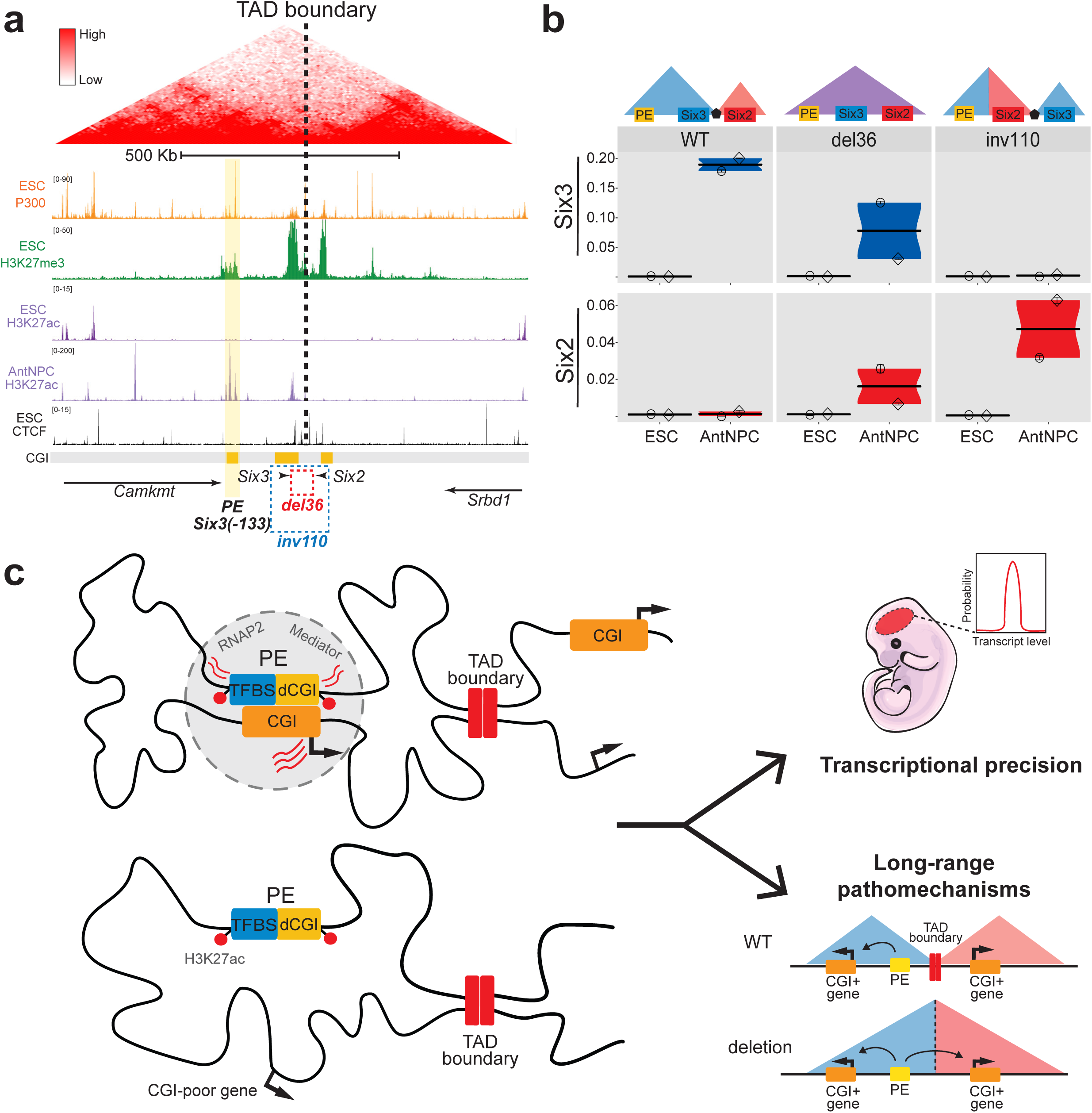
oCGI and TAD boundaries enable PEs to specifically induce their target genes. a, The TADs in which *Six3* and *Six2* are located (i.e. *Six3-*TAD and *Six2-*TAD) are shown according to publically available Hi-C data^33, 65^; the TAD boundary separating *Six3* and *Six2* is denoted with a dotted line. Below the Hi-C data, several epigenomic and genetic features of the *Six3-*TAD and the *Six2-*TAD are shown. The represented CGIs correspond to those computationally defined in the UCSC browser according to the following criteria: GC content > 50%; Length > 200bp; CpG Observed to expected ratio > 0.6. ChIP-seq profiles for the indicated proteins and histone modifications were obtained from^9, 35^. The rectangles indicate the location of the 36 Kb deletion (red) and 110 Kb inversion (blue) engineered in mESC. b, The expression of *Six3* (blue) and *Six2* (red) was measured by RT-qPCR in mESCs and AntNPC that were either WT (left panel), homozygous for the 36 Kb deletion (del36; middle panel) or homozygous for the 110 Kb inversion (inv110; right panel). For each of the engineered structural variants, two different clonal cell lines were generated and independently differentiated into AntNPC. The plotted expression values for each clone correspond to the average and standard deviation (error bars) from three RT-qPCR technical replicates. Expression values were normalized to two housekeeping genes (*Eef1a* and *Hprt*). c, Proposed model for the role of oCGI as boosters of PE regulatory activity and determinants of PE-gene compatibility. The presence of oCGI increases the physical communication of PEs with their target genes due to homotypic chromatin interactions between oCGI and promoter-proximal CGI. Consequently, the oCGI can increase the number of cells and alleles in which the PEs and their target genes are in close spatial proximity (i.e. permissive regulatory topology) both during pluripotency and upon differentiation. This will ultimately result in a timely and homogenous induction of the PE target genes once the PEs become active (i.e. increase transcriptional precision). In addition, the compatibility and responsiveness between PE and their target genes depends on the presence of oCGI and pCGI at the PEs and the target genes, respectively. Therefore, the oCGI can increase the specificity of the PEs by enabling them to preferentially communicate with their CpG-rich target genes while still being insulated by TAD boundaries. These novel PE-gene compatibility rules can improve our ability to predict and understand the pathomechanisms of human structural variants.

## Discussion

Here we show that oCGIs are an essential component of PEs that endows these distal regulatory elements with privileged regulatory properties. Namely, oCGIs boost the regulatory activity of PEs and dictate the compatibility between PEs and their target genes (Fig. 5c). Deciphering the factors that control enhancer-promoter compatibility is still a major challenge in the enhancer field that we are just starting to understand^73, 74^. Current models of enhancer function state that insulator proteins (*i.e.* CTCF) demarcate TAD boundaries and restrict enhancers to act upon genes located within their same TADs^75–78^. Nonetheless, enhancers do not promiscuously activate all the genes present within a TAD^20, 23, 79, 80^, suggesting that additional and largely unknown factors must dictate enhancer responsiveness. Using massively parallel reporter assays in *Drosophila,* it was recently shown that enhancer responsiveness is determined by the sequence composition of core promoters^25, 81^. However, we now show that, at least in the context of PE loci, such responsiveness is also dependent on distal genetic elements, namely oCGIs, which allow PEs to preferentially activate CpG rich promoters (Fig. 5c). It is worth emphasizing that neither these novel enhancer-promoter compatibility rules nor the boosting capacity of oCGIs would have been uncovered with classical reporter assays in which enhancers and core promoters are placed close to each other (Fig. 4f). Therefore, genetic engineering approaches like the one presented here will be essential to dissect the mechanisms whereby distal enhancers control the expression of their target genes.

Our data suggest that the role of oCGIs as boosters of PEs regulatory function does not involve the local activation of the PEs but rather the establishment of long-range interactions with CpG-rich gene promoters (Fig. 5c). In pluripotent cells, these PE-gene interactions are mediated by PcG complexes recruited to both oCGIs and pCGIs^8, 9, 82, 83^ and are likely to be dependent on the polymerization and/or phase separation properties of PRC1^60–62, 84^. Subsequently, PcG might keep PEs and their target genes close together during pluripotent cell differentiation, ensuring that, as the PEs become active, they can rapidly and uniformly induce the expression of their target genes. Then, once RNAs are produced at both PEs and their target genes, this would result in PcG eviction^85^. Although PRC1^63^ might also contribute to PE-gene communication once PEs become active, additional proteins and mechanisms are likely to be involved. Interestingly, we showed that, upon PE activation, the oCGI dramatically increase the loading of Mediator and the transcriptional machinery to both PEs and their CpG-rich target genes (Fig. 5c). We speculate that, by facilitating the physical communication between genes and enhancers, the oCGI might favour the formation of phase-separated transcriptional condensates^86, 87^. Once PEs are already active, multivalent interactions occurring within these condensates could robustly maintain PE-gene communication^88^.

Overall, we propose a model whereby the precise and specific induction of certain developmental genes is achieved through the combination of CGI-mediated long-range chromatin interactions and the insulation provided by TAD boundaries. We anticipate that this model may have important medical implications, as it could improve our ability to predict and understand the pathological consequences of human structural variation^12^ (Fig. 5c).

## Acknowledgements

We thank the Rada-Iglesias lab members for insightful comments and critical reading of the manuscript.

## Author Contributions

Conceptualization: T.P., A.R-I.; Experimental investigation: T.P., M.M-F., T.E., P.R., H.G.A.; Data analysis: T.P., V.S-G.; Writing, Review & Editing: T.P., A.R-I; Resources: S.C-M.,WFJ.vI., D.L., A.R.-I.; Supervision and Funding Acquisition: A.R.-I.;

## Methods

### Cell lines and differentiation protocol

mESC (E14) were cultured on gelatin-coated plates using Knock-out DMEM (KO-DMEM, Life Technologies) supplemented with 15% FBS (Life Technologies) and LIF. These mESC lines were differentiated into AntNPC according to a previously described protocol^89^, with slight modifications. Briefly, mESC were plated at a density of 12.000 cells/cm^2^ on gelatin-coated plates and grown for three days in N2B27 medium supplemented with 10 ng/ml bFGF (Life Technologies) and without serum or LIF. N2B27 medium contains: Advanced Dulbecco’s Modified Eagle Medium F12 (Life Technologies) and Neurobasal medium (Life Technologies) (1:1), supplemented with 1x N2 (Life Technologies), 1x B27 (Life Technologies), 2 mM L-glutamine (Life Technologies), 40 mg/ml BSA (Life Technologies), 0.1 mM 2-mercaptoethanol (Life Technologies)). Subsequently, cells were grown for another two days in N2B27 medium without bFgf (D3–D5). To improve the homogeneity of the differentiation, from D2-D5 the N2B27 medium was also supplemented with 5 mM Xav939, a potent WNT inhibitor^90^.

### RNA isolation, cDNA synthesis and RT-qPCR

Total RNA was isolated using Innuprep RNA mini kit (Analytik Jena) according to the manufacturer’s instructions. cDNA was generated using ProtoScript II First Strand cDNA Synthesis Kit (New England Biolabs). RT-qPCRs were performed on the Light Cycler 480II (Roche) using *Eef1a1* and *Hptr* as housekeeping genes. All primers used in RT-qPCR analysis are shown in Supplementary Data 1.

### ChIP

ChIPs were performed as previously described^8^. Basically, 50 million cells for P300/RNAP2/Med1/PcG subunits ChIPs or 10 million cells for histone ChIPs were crosslinked with 1% formaldehyde for 10 min at room temperature (RT) and then quenched with 0,125M glycine for another 10 min. Cells were then washed with PBS and resuspended sequentially in three different lysis buffers (Lysis Buffer 1: 50 mM HEPES 140 mM NaCl 1 mM EDTA 10% glycerol 0.5% NP-40 0.25% TX-100, Lysis Buffer 2: 10 mM Tris 200 mM NaCl 1 mM EDTA 0.5 mM EGTA, Lysis Buffer 3: 10 mM Tris 100 mM NaCl 1 mM EDTA 0.5 mM EGTA 0.1% Na-Deoxycholate 0.5% N-lauroylsarcosine) in order to isolate chromatin. Chromatin was then sonicated for 15 cycles (20s on 30s off, 25% amplitude) using an EpiShear probe sonicator (Active Motif). After sonication, samples were centrifuged at 16000 g during 10 min at 4°C, with the supernatant representing the sonicated chromatin. Chromatin was then incubated overnight at 4°C with 3 ug antibody for histones or 10 ug antibody for the of the investigated proteins. One of the aliquots was not subject to immunoprecipitation, thus representing total input control for the ChIP reactions. Next, 50 ul of protein G magnetic beads (Invitrogen) were added to the ChIP reactions and incubated for four additional hours at 4°C. Magnetic beads were washed and chromatin eluted, followed by reversal of the crosslinking and DNA purification. The resulting ChIP and input DNAs were analysed by qPCR using two intergenic regions as negative controls (chr2:73,030,265-73,030,373; chr6: 52,339,345-52,339,505). All primers used in ChIP-qPCR experiments are shown in Supplementary Data 1.

### Bisulfite sequencing

Bisulfite conversion of 400 ng of genomic DNA was performed using the EZ DNA Methylation Kit (Zymo Research). The *PE Sox1(+35)TFBS* or *Sox1(+35)TFBS+CGI* inserts in Gata6-TAD or *Gria1-*TAD were amplified by PCR using EpiTaq polymerase (Takara Bio) and primers described in Extended Data Fig. 7a/Supplementary Data 1. PCR products were cloned into the pGEM-T vector (Promega) and sequenced with M13 reverse primer.

### Immunofluorescence (IF)

Cells were rinsed with PBS and then fixed for 10 min in 3,7% paraformaldehyde (PFA) in PBS at RT. PFA was removed and cells were rinsed with PBS. Cells were permeabilized by treating them with 0.1% Triton X-100 for 15 minutes at RT, followed by blocking in PBS with 5%BSA for 1 hour at RT. Incubation with primary antibodies (GATA6 (AF1700, R&D systems) or SOX1 (AF3369, R&D systems)) was done in blocking solution overnight (12–18 h) at 4°C. Cells were rinsed with PBS and incubated with secondary antibodies (Life Technologies) in blocking solution for 30 minutes at RT and then rinsed again with PBS. Finally, cell nuclei were stained with DAPI (Sigma) during 10 min at RT and then mounted with anti-fading mounting medium (Life Technologies).

### 4C-seq

4C-seq Circular Chromatin Conformation Capture (4C) assays were performed as previously described^9, 91^. 1x10^7^ mESC or AntNPC were crosslinked with 1% formaldehyde during 20 minutes and quenched with 0.125M glycine for 10 minutes. Cells were washed with PBS and resuspended in lysis buffer (50 mM Tris–HCl pH 7.5, 150 mM NaCl, 5 mM EDTA, 0.5% NP-40, 1% TX-100 and 1X protease inhibitors) during 10 minutes on ice. Following centrifugation for 5 minutes at 650g at 4°C, nuclei were re-suspended in 0.5 mL of 1.2X restriction buffer with 0.3% SDS and incubated at 37°C and 900 rpm for 1 hour. After that, Triton X-100 was added to a final concentration of 2% followed by 1 hour incubation at 37°C and 900 rpm. Afterwards, chromatin was digested overnight at 37°C and 900 rpm with 400 U of NlaIII (R0125L, NEB). NlaIII was inactivated by adding SDS to a final concentration of 1.6% and incubating the mixture for 20 minutes at 65°C while shaking (900 rpm). The digested chromatin was transferred to 50 mL tubes and 6.125 mL of 1.15X ligation buffer (50 mM Tris-HCl pH 7.6, 10 mM MgCl2, 1 mM ATP, 1mM DTT) were added. Triton X-100 was also added to a final concentration of 1% and the resulting solution was incubated for 1 hour at 37°C while shaking gently. After that, digested chromatin was ligated with 100 U of T4 DNA ligase (15224-041, Life Technologies) for 8 hours at 16°C, followed by RNase A treatment (Peqlab) for 45 minutes at 37°C. Subsequently, chromatin was de-crosslinked with 300 mg of Proteinase K (Peqlab) and incubated at 65°C overnight. DNA was then purified by phenol/chloroform extraction followed by ethanol precipitation and re-suspension in 100 mL of water. At this step, the digestion and ligation efficiencies were evaluated by analysing a small fraction of the purified DNAs by agarose electrophoresis. The remaining DNA was digested with 50U of DpnII (R0543M, NEB) at 37°C overnight. DNA samples were then purified by phenol/chlorophorm extraction, followed by ethanol precipitation and resuspension in 500 ul of H2O. Afterwards, a second ligation was performed by adding 200 U of T4 DNA Ligase into a final volume of 14 mL 1X Ligation Buffer and incubating overnight at 16°C. DNA samples were subjected to another round of phenol/chlorophorm extraction and ethanol precipitation, re-suspended in 100 uL of water and purified with a QIAgen PCR purification column (28104, QIAgen). Finally, the resulting 4C-DNA products were amplified by inverse PCR using primers located within selected PEs, which were designed as previously described^91^ (Supplementary Data 1). The inverse PCRs were performed with the expand long template PCR system (11681842001, Roche) using 30 amplification cycles (94°C 2 min, 30x [94°C 10s,60°C 1 min, 68°C 3 min], 68°C 5 min). 4C-seq libraries were then analysed by next generation sequencing. All the generated 4C-seq data will be deposited in GEO and made available upon publication.

### oCGI deletion using CRISPR-Cas9

To generate the deletion of the *PE Sox1(+35)CGI*, pairs of sgRNAs flanking the oCGI were selected according to Benchlinǵs CRISPR toolkit (www.benchling.com) (Supplementary Data 1). For each selected sgRNA, two oligonucleotides were synthesized (Integrated DNA Technologies) and annealed. The CRISPR-Cas9 expression vector, pX330-hCas9-long-chimeric-grna-g2p (provided by Leo Kurian’s laboratory), was digested with BbsI (R0539L, NEB) and gel or column purified. Pairs of annealed oligos and the digested vector were ligated overnight at 16°C using T4 ligase (NEB). Following transformation, the gRNA-Cas9 expression vectors were purified and sequenced to confirm that the gRNAs were correctly cloned. mESC were transfected with the pair of gRNAs-Cas9 expressing vectors using Lipofectamine according to the manufacturer protocol (Thermo Scientific). After 16 hours, puromycin selection was performed for 48 hours. Subsequently, surviving cells were isolated in 96-well plates by serial dilution and, following expansion, clones with the desired deletion were identified by PCR using the primers listed in Supplementary Data 1. Finally, the presence of the desired deletion was confirmed in the selected mESC clones by Sanger sequencing.

### Homology-dependent Knock-in

Knock-In of PE modules was performed using CRISPR-Cas9 as previously described by Yao and colleagues^32^ with minor modifications. First, a sgRNA was designed for the insertion site of interest and cloned in a CRISPR-Cas9 expression vector as mentioned above. Then, the *cassette-vector* was generated by ligating: (i) 300bp homology arms flanking the insertion site; (ii) construct of interest; and (iii) cloning vector. The resulting *cassette-vector* was used as a template for amplifying the *knock-in donor* (left homology arm + construct + right homology arm) by PCR (Supplementary Data 2). The resulting PCR product was purified using QIAgen PCR purification columns (28104, QIAgen). mESC were transfected with the sgRNA-Cas9 expressing vector and the *knock-in donor* using Lipofectamine according to the manufacturer protocol (Thermo Scientific). After 16 hours, puromycin selection was performed for 48 hours. Subsequently, surviving cells were isolated in 96-well plates by serial dilution and, following expansion, clones with the desired insertions were identified by PCR using the primers listed in Supplementary Data 1. Finally, the insertion of the desired PE modules was confirmed in the selected mESC clones by Sanger sequencing.

### FAIRE (Formaldehyde-Assisted Isolation of Regulatory Elements)

Sonicated chromatin was prepared as for ChIP and then DNA was purified as previously described^92^. Briefly, chromatin was subject to three rounds of phenol/chloroform purification and the resulting DNA was purified by precipitation with sodium acetate and ethanol. The resulting FAIRE and input DNAs were analysed by qPCR using two intergenic regions as negative controls (chr2:73,030,265-73,030,373; chr6:52,339,345-52,339,505). All primers used in the FAIRE-qPCR experiment are shown in Supplementary Data 1.

### NOMe-PCR

Nuclei extraction and M.CviPI treatment were performed as described previously^93^ with minor modifications. Basically, isolated nuclei were incubated with 200 U of M.CviPI (NEB) for 15 min at 37 °C. Then, bisulfite conversion was performed using the EZ DNA Methylation Kit (Zymo Research) and the converted DNA was amplified by PCR. Finally, the PCR product were cloned into the pGEM-T vector (Promega) and sequenced with M13 reverse primer. NOMe-PCR data was analysed with the NOMePlot web app tool (http://www.landeiralab.ugr.es/software)^94^.

## Computational and Statistical Analyses

### Analyses of q-PCR data

For RT-qPCR, relative gene expression levels were calculated using the 2^ΔCt^ method. Standard deviations were calculated from technical triplicate reactions and were represented as error bars. Primers used can be found in Supplementary Data 1.

ChIP-qPCR signals were calculated as % of input using technical triplicates. Each ChIP sample was normalized to the average signals obtained in the same sample when using negative control regions primers (Chr2_neg and Chr6_neg; see Supplementary Data 1). Standard deviations were calculated from technical triplicate reactions and represented as error bars.

### 4C-seq analysis

All 4C-seq samples were sequenced on an Illumina HiSeq 2500 sequencer, generating single reads of 74 bases in length. From these reads, the sequence attached to the viewpoint was extracted starting before the restriction site for NlaIII (CATG). These sequences were trimmed down to 41 base-pairs and aligned to the mouse GRCm38/mm10 reference genome using the HISAT2 aligner^87^. From these alignments, RPM (reads per million) normalized bedgraph files were generated for downstream visualization and analysis^88^.

### Gene Annotation

The RefSeq gene annotation was downloaded through the UCSC Table Browser (https://genome.ucsc.edu/cgi-bin/hgTables) and used for the different analyses,

### ChIP-Seq and PRO-Seq pre-processing steps

For all ChIP-Seq or PRO-Seq samples analyzed from fastq reads, the read quality was assessed with FastQC (http://www.bioinformatics.babraham.ac.uk/projects/fastqc/) and MultiQC^95^.

For ChIP-Seq data, the removal of read adapters and low quality filtering was done with trimmomatic^96^.

For PRO-Seq data, adapter removal was performed with cutadapt 1.18^97^ filtering for a minimum of 15 bases (adapter sequence: TGGAATTCTCGGGTGCCAAGG). In addition, reads mapping to the mouse rDNA repeats (GenBank: BK000964.3) were discarded from downstream analysis.

For both data types, reads were mapped to the mouse mm9 reference genome with Bowtie2^98^. Last, for all ChIP-Seq samples, upon read mapping, duplicated reads were discarded, with the usage of SAMtools^99^.

### Genetic properties of CGIs

#### Data retrieval and pre-processing

To perform the analyses described below, the required data sets were: PE coordinates, CGI coordinates and a list of developmental genes.

Poised enhancer coordinates were downloaded from ^9^ and converted from *mm10* to *mm9* mouse genome coordinate with the UCSC LiftOver tool (https://genome.ucsc.edu/cgi-bin/hgLiftOver<u>). Only poised enhancers more than 2.5kb away from any TSS (PE-all) were considered. Among them, those PEs that were at least 10kb away from any TSS are referred to as *PE-distal*. All RefSeq transcripts were considered for this PE classification.</u>

NMI coordinates were obtained from ^28^. CAP-CGI coordinates were obtained from ^7^, without applying any extension.

A relevant feature for many important cell-identity genes is that they are frequently embedded within broad domains of H3K27me3 when they are not expressed^100, 101^. Therefore, we processed H3K27me3 ChIP-Seq and its corresponding input data generated in mESC (GSE89209; H3K27me3 ID: SRR4453259, Input ID: SRR4453262). Next, H3K27me3 peaks with respect to the Input were called with MACS2^102^ using the broad peak calling mode. Only those peaks with a fold-enrichment > 3 and q value < 0.1 were maintained. Subsequently, peaks closer than 1kb were merged using bedtools, and associated with a protein coding gene if they overlapped a TSS. Lastly, the size distribution of the H3K27me3 peaks associated with genes was studied and developmental genes were defined as those genes with the largest peak sizes. To do so, the knee of the size distribution was determined with *findiplist()* (*inflection* R package; [https://cran.r-project.org/web/packages/inflection/vignettes/inflection.html]). Upon curvature analysis, a threshold of 6 Kb was defined and, thus, all genes with a H3K27me3 peak length larger than 6kb were considered as developmental genes (devTSS).

#### CGI groups classification

NMI and CAP-CGI were associated with PE-distal or devTSS if located less than 3kb away from them. In addition, to get a representation of the bulk genome composition, a third group of random regions was created (generated independently for NMI and CAP-CGI sequence composition analysis). To create this random group, the set of regions (NMI or CAP-CGI) in proximity to PE-distal was taken as reference in terms of region sizes. Then, each of the regions associated with a PE-distal was randomly relocated along the genome 1000 times (maintaining its size). Finally the set of all randomized regions constituted the random group.

#### Sequence Composition Calculations

To retrieve the DNA sequences of the studied regions, the BSgenome package was used^103^, using as reference the unmasked mouse mm9 genome. For each region: its length, the percentage in G+C, the percentage in CpG and the CpG observed/expected ratio was calculated. The %CpG was calculated as the ratio of CpG dinucleotide counts with respect to half the total region length. The ratio of observed to expected CpG is calculated according to the formula presented in ^104^.

### Comparison of eRNA levels between different classes of active enhancers

#### Data retrieval and pre-processing

The required data sets were: active enhancer coordinates, TAD maps, RNA-Seq, H3K27ac and PRO-Seq data from WT mESC. The active enhancer coordinates and the RNA-Seq data were used in both of the analyses described below.

Gene expression data (RNA-Seq FPKMs) and active enhancer coordinates from WT mESC were obtained from ^9^. Enhancer coordinates were converted from mm10 to mm9 genome version with the UCSC LiftOver tool (https://genome.ucsc.edu/cgi-bin/hgLiftOver). To avoid confounding effects between transcripts produced by enhancers or genes, only active intergenic enhancers located at least 10kb away from any TSS and 20kb from any transcription termination site (TTS) were considered.

Regarding the H3K27ac ChIP-Seq data from mESC, for the analyses presented in Fig 4 and Extended Fig 11a, the fastq raw data was retrieved from GEO (GSE89209; sample ID: <u>SRR4453258</u>) and pre-processed as indicated above. For the analyses presented in Extended Fig 11b-c, two H3K27ac bigWig files (from two replicates) were downloaded from GEO (samples IDs: GSM2808655 and GSM2808669).

With respect to the PRO-Seq data from mESC, for the analyses presented in Fig 4 and Extended Fig 11a, the fastq raw data was obtained from GEO (GSE115713; sample IDs: <u>SRR7300121</u>, <u>SRR7300122</u>). The data from the two replicates was combined and pre-processed as described above. For the analyses presented in Extended Fig 11b-c, two PRO-Seq bigWig files (one for the forward DNA strand and the other for the reverse one) derived from the combination of three replicates, were obtained from GEO (GSE130691).

TAD maps from mESC were retrieved from ^33^. For the analyses presented in Fig 4 and Extended Fig 11a, the TAD map used was mESC_Dixon2012-raw_TADs.txt. For the analyses presented in Extended Fig 11b-c, the TAD map used was mESC.Bonev_2017-raw.domains, whose coordinates were converted from mm10 to mm9 with UCSC LiftOver tool (https://genome.ucsc.edu/cgi-bin/hgLiftOver).

#### H3K27ac & PRO-Seq enhancer levels quantification

To quantify H3K27ac and PRO-Seq enhancer levels two different approaches were followed, one for the analyses presented in Fig 4 and Extended Fig 11a .and another one for the analyses presented in Extended Fig 11b-c.

Fig 4 and Extended Fig 11a: H3K27ac and PRO-Seq reads presenting a mapping quality less than 10 were discarded using SAMtools^99^. Next, with the resulting mapped reads in BAM format, a bigwig was generated with *deepTools*^105^ and *bamCoverage* tool (RPGC normalization, and 1870000000 effective genome size). Finally, H3K27ac and PRO-Seq enhancer mean scores were obtained with the computeMatrix tool from *deepTools*, taking as input the active enhancer coordinates and the H3K27ac and PRO-Seq bigwigs. For H3K27ac, the signals were calculated +/-1kb with respect to the enhancer midpoints, while for PRO-Seq a +/-0.5kb window was used instead.

Extended Fig 11b-c: H3K27ac and PRO-Seq mean signals for the active enhancers were calculated with the bigWigAverageOverBed UCSC binary tool. Next, PRO-Seq signals for each enhancer from the two different strands were averaged and the same was done for the signals coming from the different H3K27ac replicates.

#### Active enhancers classification

Three groups of active enhancers were defined according to the following criteria. (*I*) enhancers located in TADs containing only poorly expressed genes (all genes with <0.5 FPKM); (*II*) enhancers located in a TAD with at least one gene with expression levels >10 FPKM; (*III*) enhancers whose closest gene within the same TAD has expression levels >10 FPKM.

#### Balancing of H3K27ac levels within enhancer classes

To evaluate whether differences in eRNA levels between the three groups of active enhancers could be simply explained by differences in H3K27ac, the levels of this histone modification were balanced in some of the presented analyses (Extended Fig 11a,c). Briefly, enhancers with similar H3K27ac levels belonging to the three enhancer classes were selected by applying the nearest neighbour matching method (without replacement and ratio = 1) using *MatchIt* [https://cran.r-project.org/web/packages/MatchIt/MatchIt.pdf] and considering the enhancer group (I) as the treatment condition.

#### Cliff’s delta effect size estimator

Cliff’s delta^106, 107^, a non-parametric effect size estimator, was used to quantify and interpret the differences between different groups of genomic regions for different types of genetic, transcriptional or epigenetic features. This measure relies on the concept of dominance (which refers to the degree of overlap between distributions) rather than means (as in conventional effect size indices such as Cohen’s *d*) and is considered to be more robust when signal distributions are skewed^108^. Cliff’s delta, was estimated using the cliff.delta() function from the R package *effsize* [https://cran.r-project.org/web/packages/effsize/index.html]. Differences between groups with an associated |delta| <0.147 can be considered as negligible and |delta|>= 0.147 as non-negligible.

---

**Extended Data Fig. 1.**
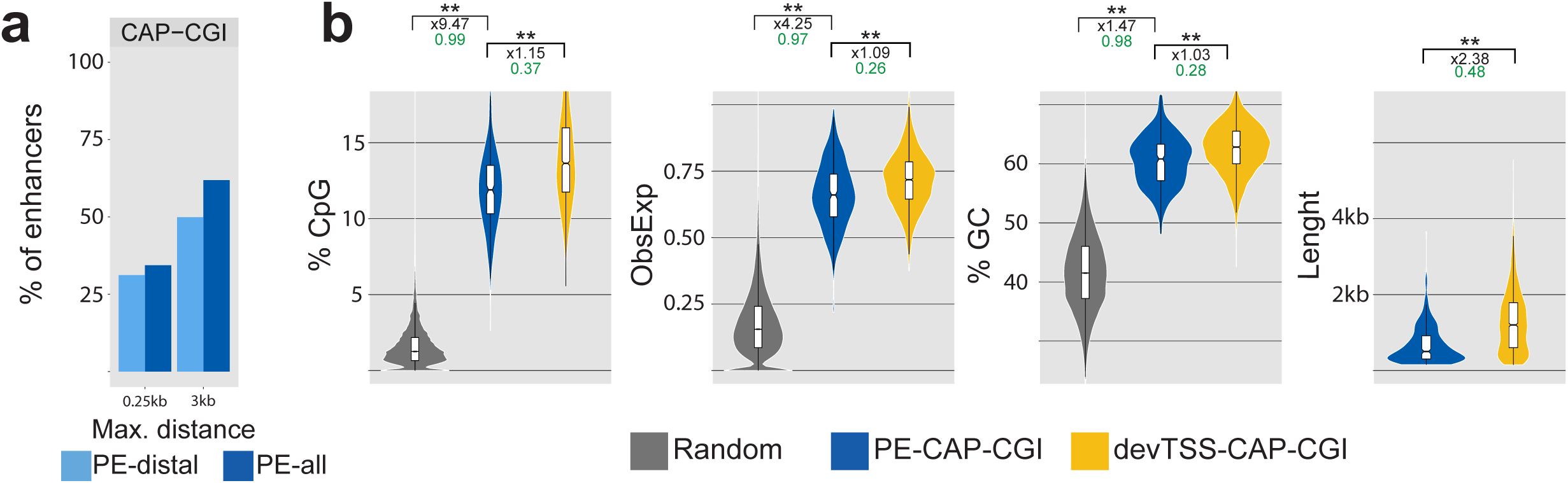
Genetic features of the orphan CGIs associated with PEs. a, Percentage of poised enhancers (PEs) found within the indicated maximum distances (0.25Kb or 3Kb) to a computationally-defined CGI according to the following criteria: GC content > 50%; Length > 200 bp; CpG (left panel) or a CAP-CGI identified with the CAP assay^7^ (right panel). b, Comparison of the CpG percentage, observed/expected CpG ratio, GC percentage and sequence length between random regions (see Methods), CAP-CGIs associated to poised enhancers (PE-CAP-CGI; blue) and CAP-CGIs associated to the TSS of developmental genes (devTSS-CAP-CGI; yellow; see Methods). On top of each plot, the asterisks indicate P-values calculated using unpaired Wilcoxon tests with Bonferroni correction for multiple testing (** = p.val < 1e^-10^; * p.val < 0.05); the numbers in black indicate the median fold changes between the indicated groups; the green numbers indicate non-negligible Cliff Delta effect sizes.

**Extended Data Fig. 2.**
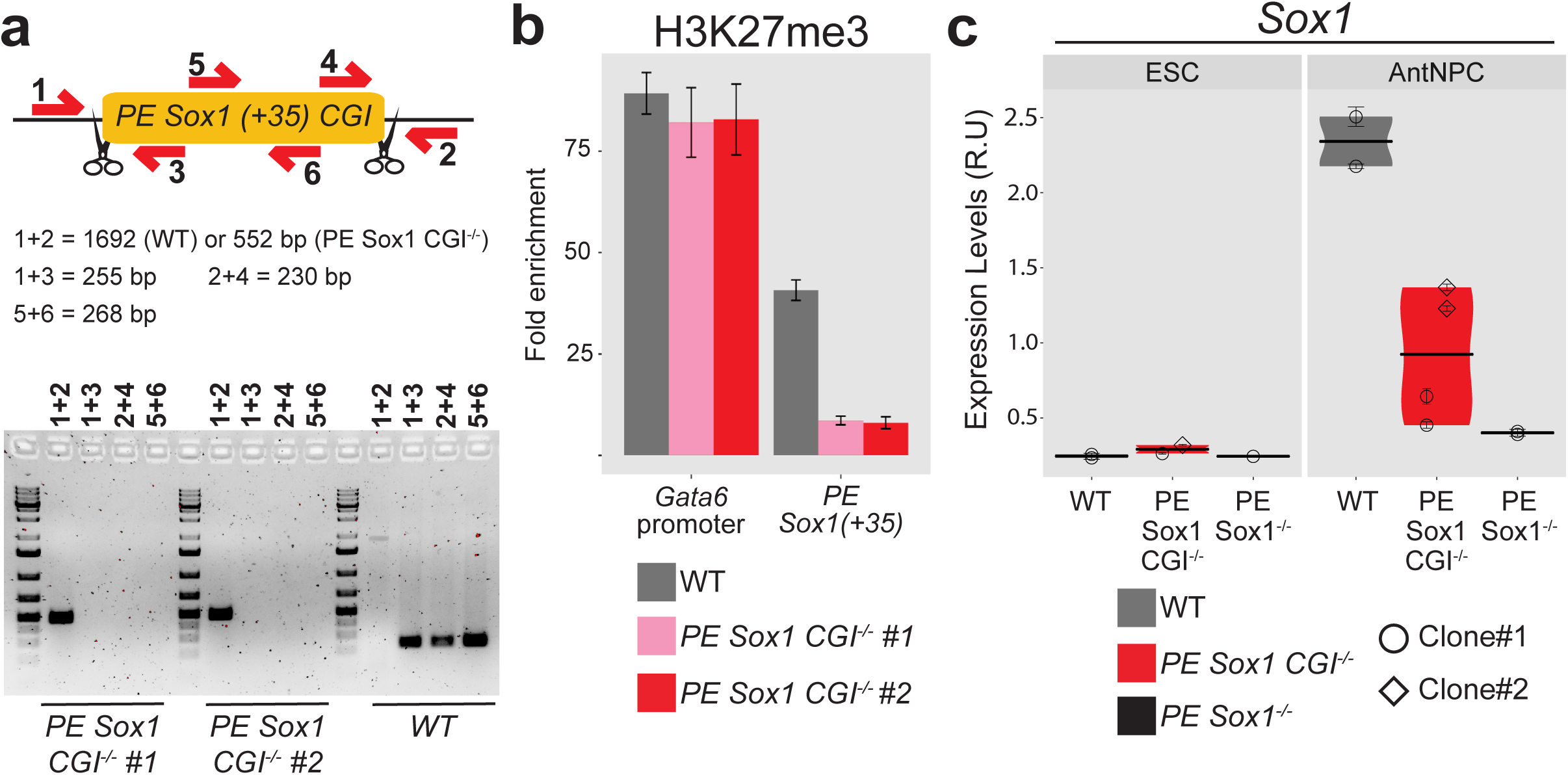
*PE Sox1(+35)CGI* recruits PcG and contributes to the cis-regulatory function of *PE Sox1(+35)*. a, For the identification of the *PE Sox1(+35)CGI* deletion, primer pairs flanking each of the deletion breakpoints (1+3 and 4+2), located within the deleted region (5+6) or amplifying a large or small fragment depending on the absence or presence of the deletion (1+2) were used. The PCR results obtained for WT ESC and for two mESC clonal lines with homozygous deletions of the PE *Sox1(+35)CGI* (*PE Sox1(+35)CGI^-/^*) are shown. b, H3K27me3 levels at *PE Sox1(+35)* were measured by ChIP-qPCR in WT mESC (grey), and in two *PE Sox1(+35)CG^I-/-^* mESCs clones using primers adjacent to the deleted region. ChIP-qPCR signals were normalized against two negative regions (Supplementary Data 1). Error bars correspond to standard deviations from technical triplicates. c, Independent biological replicate for the data presented in Fig. 1d. The expression of *Sox1* was investigated by RT-qPCR in mESCs (left panel) and AntNPC (right panel) that were either WT (grey), homozygous for a deletion of the *PE Sox1(+35) CGI* (*PE Sox1 CGI^-/-^*; red) or homozygous for the complete *PE Sox1(+35)* deletion^9^ (*PE Sox1^-/-^*; black). Two different *PE Sox1 CGI^-/-^* mESC clones (circles and diamonds) and one *PE Sox1^-/-^* clone were studied. For each cell line, two technical replicates of the AntNPC differentiation were performed. The plotted expression values of each clone correspond to the average and standard deviation (error bars) from three RT-qPCR technical replicates. Expression values were normalized to two housekeeping genes (*Eef1a* and *Hprt*).

**Extended Data Fig.3.**
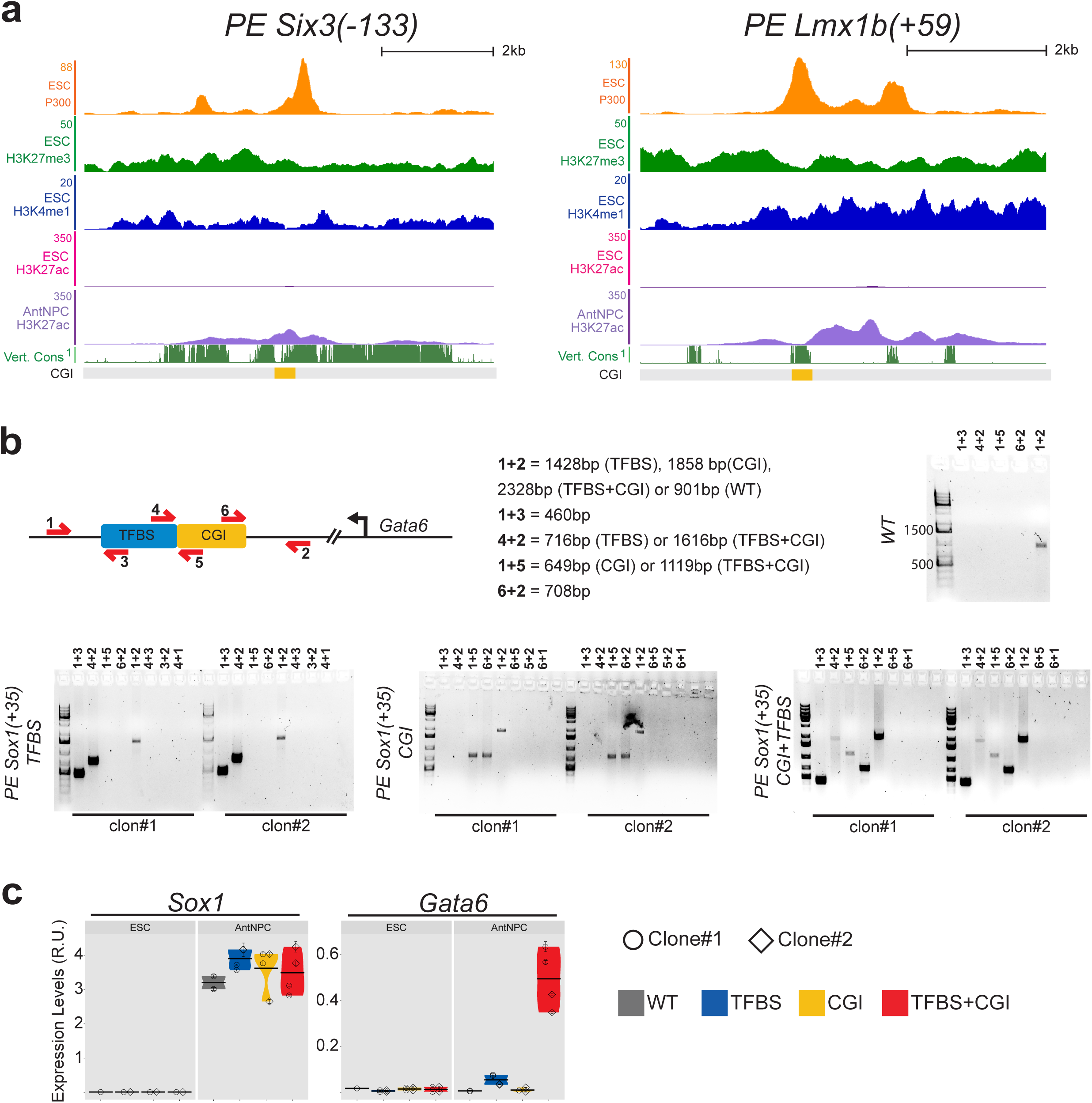
Modular engineering of the *PE Sox1(+35)* within the *Gata6*-TAD. a, Genome-browser view of the epigenomic and genomic features of two previously characterized PEs^9^ (left: *PE Six3-(133)*; Right: *PE Lmx1b(+59)*) in which the oCGIs overlap with conserved sequences bound by P300 and, thus, likely to represent TFBS. The represented CGIs correspond to those computationally defined in the UCSC browser according to the following criteria: GC content > 50%; Length > 200 bp; CpG Observed to expected ratio > 0.6. Vert. Cons.= vertebrate PhastCons. b, For the identification of the different *PE Sox1(+35)* module insertions, primer pairs flanking the insertion borders (1+3 and 4+2; 1+5 and 6+2; or 1+3 and 6+2), amplifying potential duplications (4+3, 3+2 and 4+1; or 6+5, 5+2 and 6+1) and amplifying a large or small fragment depending on the absence or presence of the insertion (1+2), respectively, were used. The PCR results obtained for WT mESC and for two mESC clonal lines with homozygous insertions for each of the three different combinations of *PE Sox1(+35)* modules (i.e. (i) *PE Sox1(+35)TFBS*; (ii) *PE Sox1(+35)CGI*; (iii) *PE Sox1(+35)TFBS+CGI*) inserted in the *Gata6-*TAD are shown. c, Independent biological replicate for the data presented in Fig. 2b. The expression of *Gata6* and *Sox1* was measured by RT-qPCR in mESCs (left panels) and AntNPC (right panels) that were either WT (grey) or homozygous for the insertions of the different *PE Sox1(+35)* modules (i.e. TFBS (blue), CGI (yellow), TFBS+CGI (red)). For the cells with the PE module insertions, two different clonal cell lines (circles and diamonds) were studied in each case. For each cell line, two technical replicates of the AntNPC differentiation were performed. The plotted expression values for each clone correspond to the average and standard deviation (error bars) from three RT-qPCR technical replicates. Expression values were normalized to two housekeeping genes (*Eef1a* and *Hprt*).

**Extended Data Fig.4.**
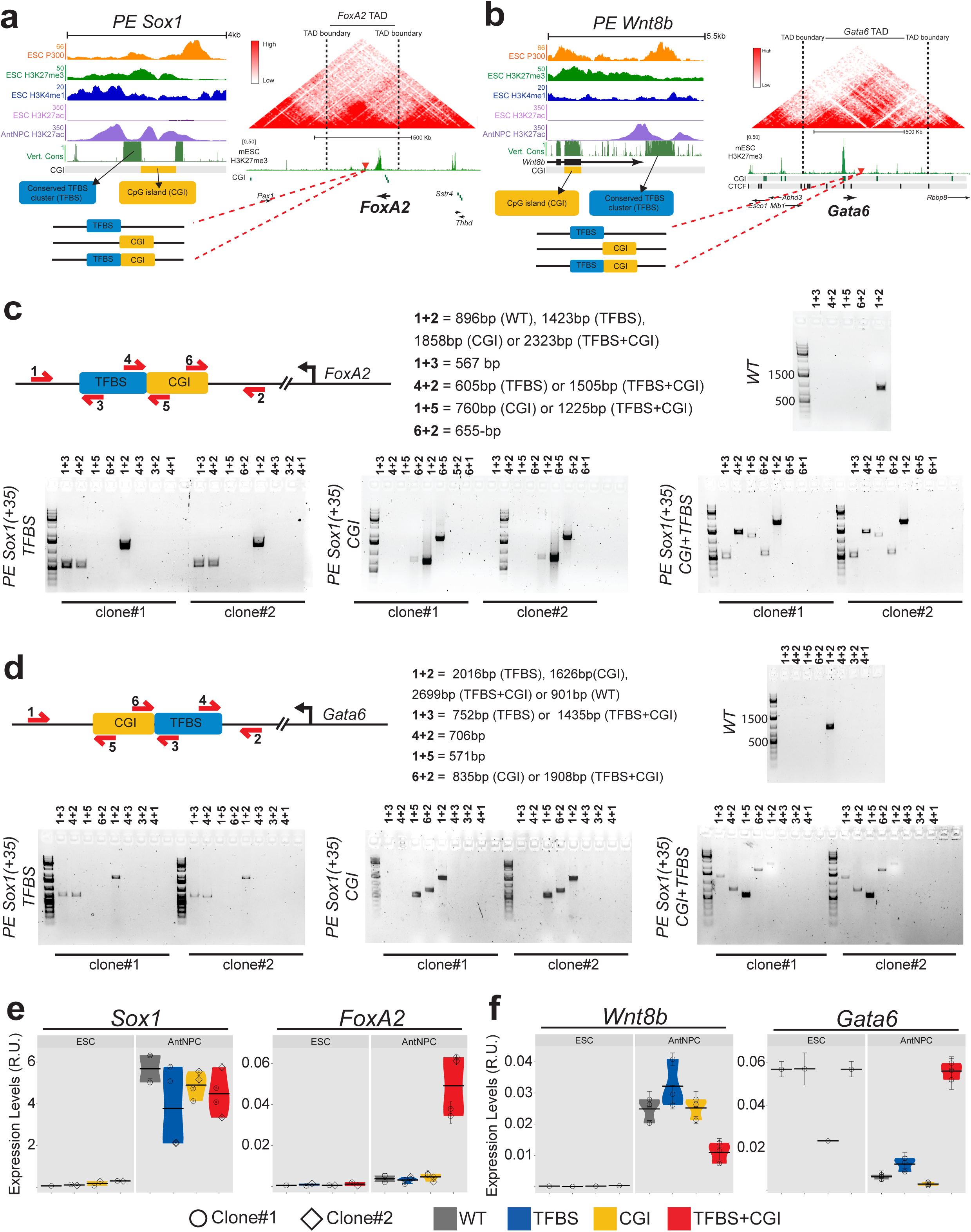
Modular engineering of the *PE Sox1(+35)* within the *FoxA2*-TAD and of the *PE Wnt8b(+21)* within the *Gata6*-TAD. a, Strategy used to insert the *PE Sox1(+35)* components into the *Foxa2*-TAD. The upper left panel shows a close-up view of the epigenomic and genetic features of the *PE Sox1(+35)* (Vert. Cons.= vertebrate PhastCons). The represented CGIs correspond to those computationally defined in the UCSC browser according to the following criteria: GC content > 50%; Length > 200 bp; CpG Observed to expected ratio > 0.6. The lower left panel shows the three combinations of *PE Sox1(+35)* modules (i.e. (i) *PE Sox1(+35)TFBS*; (ii) *PE Sox1(+35)CGI*; (iii) *PE Sox1(+35)TFBS+CGI*) inserted into the *FoxA2*-TAD. The right panel shows the TAD in which *Foxa2* is included (i.e. *Foxa2*-TAD) according to publically available Hi-C data^33, 65^; TAD boundaries are denoted with dotted lines; H3K27me3 ChIP-seq signals in mESC are shown in green^9^; CGIs are indicated as green rectangles; the red triangle indicates the integration site of the *PE Sox1(+35)* modules, approximately 100 Kb downstream of *Foxa2*. b, Strategy used to insert the *PE Wnt8b(+21)* components into the *Gata6-*TAD as described in (a). c-d, For identifying the successful insertion of the different *PE Sox1(+35)* (c) or *PE Wnt8b(+21)* (d) modules, primer pairs flanking the insertion borders (1+3 and 4+2; 1+5 and 6+2; or 1+3 and 6+2), amplifying potential duplications (4+3, 3+2 and 4+1; or 6+5, 5+2 and 6+1) and amplifying a large or small fragment depending on the absence or presence of the insertion (1+2), respectively, were used. The PCR results obtained for two mESC clonal lines with homozygous insertions for each of the three different combinations of *PE Sox1(+35)* modules (i.e. (i) *PE Sox1(+35)TFBS*; (ii) *PE Sox1(+35)CGI*; (iii) *PE Sox1(+35)TFBS+CGI*) or *PE Wnt8b(+21)* modules (i.e. (i) *PE Wnt8b(+21)TFBS*; (ii) *PE Wnt8b(+21)CGI*; (iii) *PE Wnt8b(+21)TFBS+CGI*) in the *Foxa2-*TAD (c) or *Gata6*-TAD (d), respectively, are shown. e-f, Independent biological replicate for the data shown in Fig. 2c (e) and Fig. 2d (f). The expression of *Foxa2* (e), *Gata6* (f), *Sox1* (e) and *Wnt8b* (f) was measured by RT-qPCR in mESCs (left panels) and AntNPC (right panels) that were either WT (grey) or homozygous for the insertions of the different PE *Sox1(+35)* (e) or *Wnt8b(+21)* (e) modules (i.e. TFBS (blue), CGI (yellow), TFBS+CGI (red)). For the cells with the PE module insertions, two different clonal cell lines (circles and diamonds) were studied in each case. For each cell line, two technical replicates of the AntNPC differentiation were performed. The plotted expression values for each clone correspond to the average and standard deviation (error bars) from three RT-qPCR technical replicates. Expression values were normalized to two house-keeping genes (*Eef1a* and *Hprt*).

**Extended Data Fig.5.**
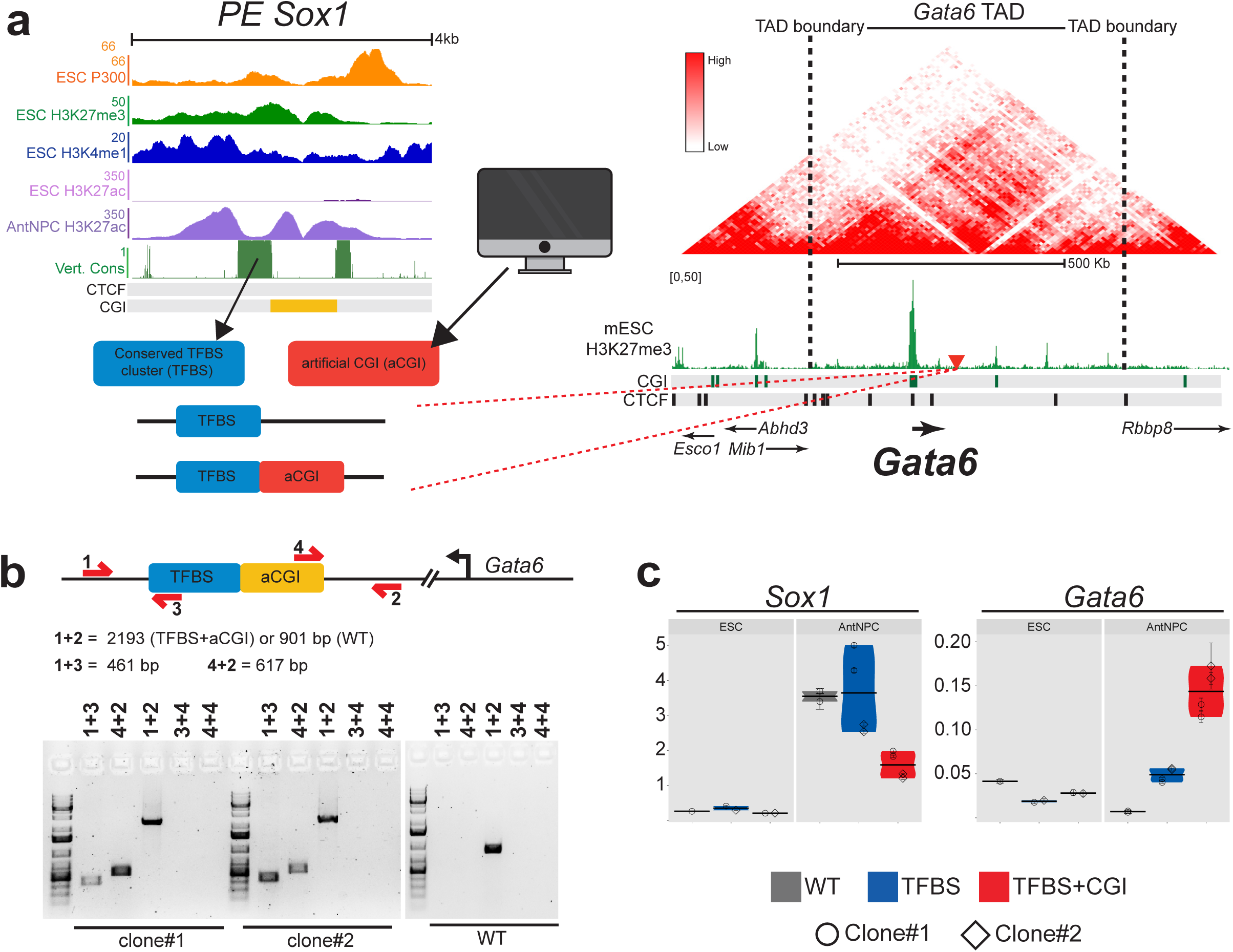
Engineering of a *PE Sox1(+35*) construct with the TFBS module and an artificial CGI. a, Strategy used to insert the *PE Sox1(+35)TFBS* alone of together with an artificial CGI (aCGI; see Methods) into the *Gata6*-TAD. The upper left panel shows a close-up view of the epigenomic and genetic features of the *PE Sox1(+35)* (Vert. Cons.= vertebrate PhastCons). The represented CGIs correspond to those computationally defined in the UCSC browser according to the following criteria: GC content > 50%; Length > 200 bp; CpG Observed to expected ratio > 0.6. The lower left panel shows the two combinations of *PE Sox1(+35)* modules (i.e. (i) *PE Sox1(+35)TFBS*; (ii) *PE Sox1(+35)TFBS+aCGI*) inserted into the *Gata6* TAD. The right panel shows the TAD in which *Gata6* is included (i.e. *Gata6*-TAD) according to publically available Hi-C data^33, 34^; TAD boundaries are denoted with dotted lines; H3K27me3 ChIP-seq signals in mESC are shown in green^9^; CGIs are indicated as green rectangles; CTCF binding sites^35^ are indicated as black rectangles; the red triangle indicates the integration site of the *PE Sox1(+35)* modules, approximately 100 Kb downstream of *Gata6*. b, For the identification of the *PE Sox1(+35)TFBS+aCGI* insertion, primer pairs flanking the insertion borders (1+3 and 4+2), amplifying potential duplications (4+3 and 4+4) and amplifying a large or small fragment depending on the absence or presence of the insertion (1+2), respectively, were used. The PCR results obtained for two mESC clonal lines with homozygous insertions of *PE Sox1(+35)TFBS+aCGI* in the *Gata6-*TAD are shown. c, Independent biological replicate for the data presented in Fig. 2e. The expression of *Gata6* and *Sox1* was measured by RT-qPCR in mESCs (left panels) and AntNPC (right panels) that were either WT (grey) or homozygous for the *PE Sox1(+35)TFBS* (blue) or *PE Sox1(+35)TFBS+aCGI* (red) insertions. For the cells with the PE insertions, two different clonal cell lines (circles and diamonds) were studied in each case. For each cell line, two technical replicates of the AntNPC differentiation were performed. The plotted expression values for each clone correspond to the average and standard deviation (error bars) from three RT-qPCR technical replicates. Expression values were normalized to two house-keeping genes (*Eef1a* and *Hprt*).

**Extended Data Fig.6.**
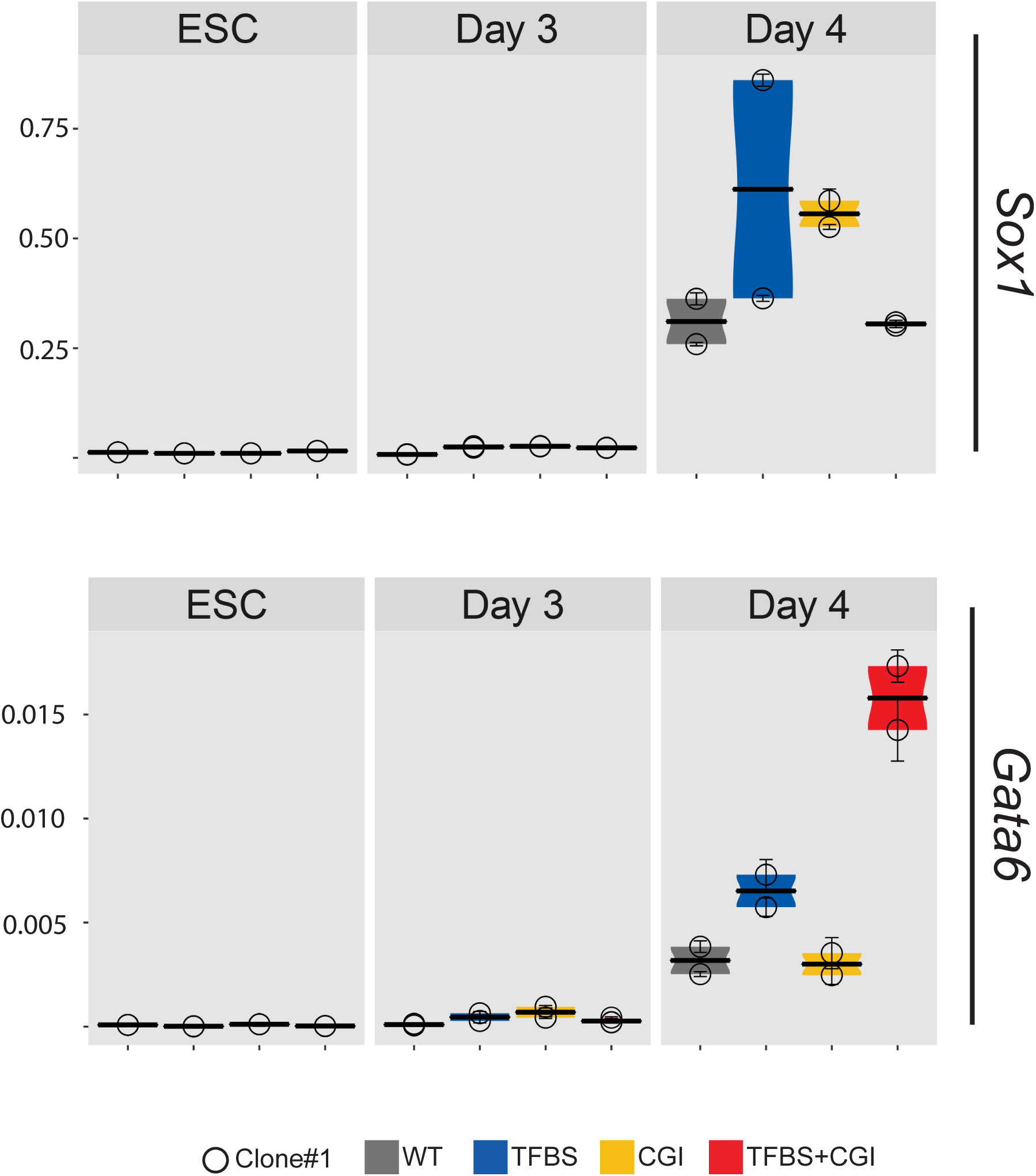
*Gata6* expression dynamics in cell lines with the *PE Sox1(+35)* modules inserted within the *Gata6*-TAD. The expression of *Gata6* and *Sox1* was measured by RT-qPCR in mESCs and at intermediate stages of mESCs differentiation into AntNPC (Day 3 and Day 4). The analysed cells were either WT (grey) or homozygous for the insertions of the different *PE Sox1(+35)* modules (i.e. TFBS (blue), CGI (yellow), TFBS+CGI (red)) within the *Gata6-*TAD. For the cells with the PE module insertions, one clonal cell line (circles) was studied. For each cell line, two technical replicates of the AntNPC differentiation were performed. The plotted expression values for each clone correspond to the average and standard deviation (error bars) from three RT-qPCR technical replicates. Expression values were normalized to two housekeeping genes (*Eef1a* and *Hprt*).

**Extended Data Fig.7.**
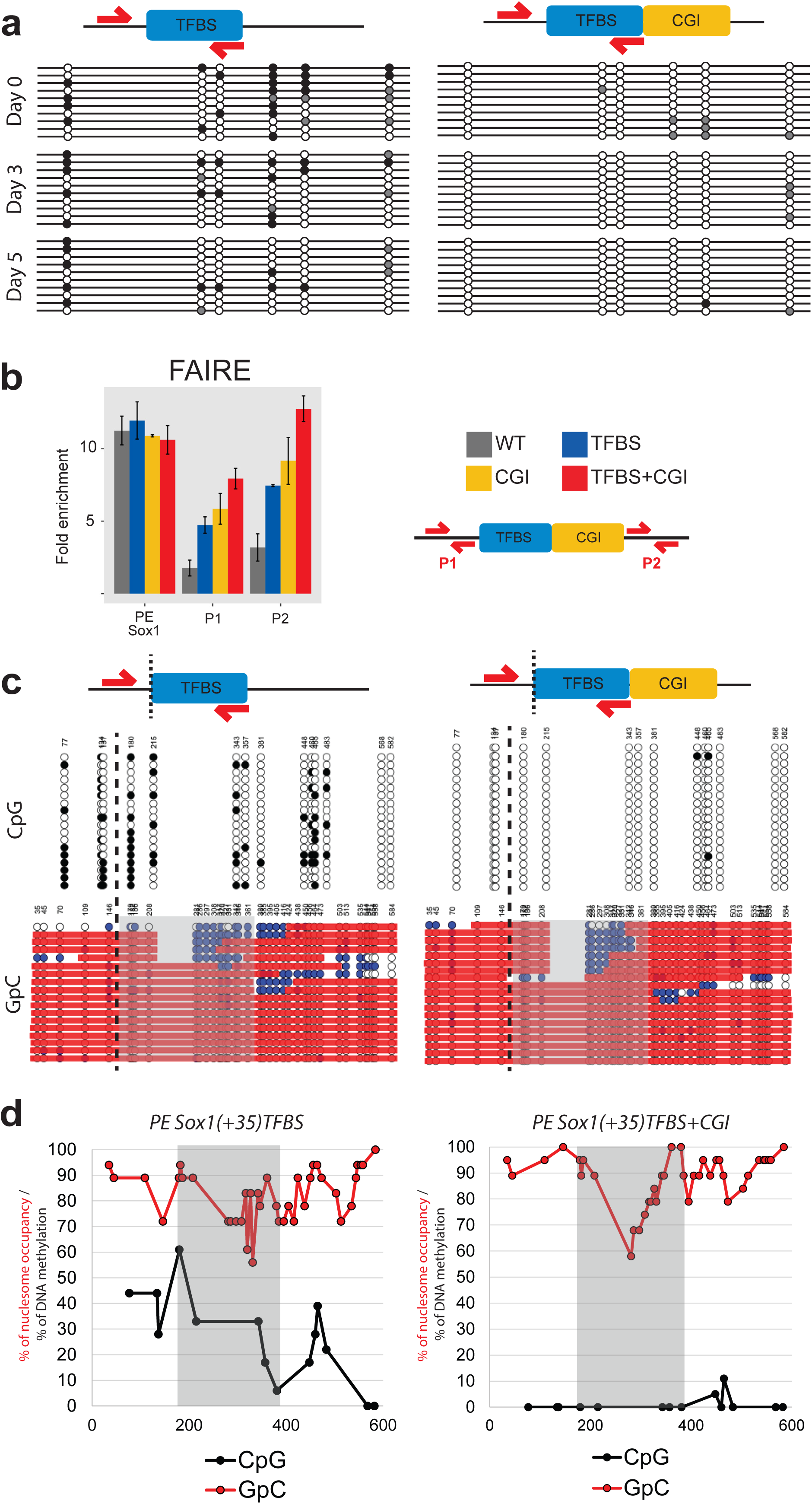
Epigenetic characterization of the PE Sox1(+35) modules engineered within the Gata6-TAD. **a**, Detailed view of the bisulfite sequencing data presented in Fig. 3a, in which mESC (Day0) and AntNPC (Day5) differentiated from cell lines with the PE *Sox1(+35)TFBS* (left panel) or *PE Sox1(+35)TFBS+CGI* modules (right panel) inserted in the *Gata6-*TAD were analyzed. DNA methylation levels were measured using a forward bisulfite primer upstream of the insertion site and a reverse primer inside the TFBS module (Methods). The circles in the plots correspond to individual CpG dinucleotides located within the TFBS module. Unmethylated CpGs are shown in white, methylated CpGs in black and not-covered CpGs are shown in gray. 10 alleles (rows) were analyzed for each differentiation stage and cell line. **b**, Chromatin accessibility at the endogenous *PE Sox1(+35)*, the *Gata6* TAD insertion site (primer pairs P1 and P2) and the *Gata6* promoter were measured by FAIRE-qPCR in mESCs (left panels) and AntNPC (right panels) that were either WT (gray) or homozygous for the insertions of the different *PE Sox1(+35)* modules (i.e. TFBS (blue), CGI (yellow), TFBS+CGI (red)). FAIRE-qPCR signals were normalized against two negative control regions (Supplementary Data 1). Error bars correspond to standard deviations from technical triplicates. The location of the primer pairs P1 and P2 around the *Gata6-*TAD insertion site is represented as red arrows in the diagram shown to the right. **c**, DNA methylation and nucleosome occupancy at the TFBS module were simultaneously analyzed by NOME-PCR in ESC lines with the PE *Sox1(+35)TFBS* (left panel) or *PE Sox1(+35)TFBS+CGI* modules (right panel) inserted in the *Gata6-*TAD. In the upper panels, the black and white circles represent methylated or unmethylated CpG sites, respectively. In the lower panels, the blue or white circles represent accessible or inaccessible GpC sites for the GpC methyltransferase, respectively. Red bars represent regions large enough to accommodate a nucleosome and that are considered as inaccessible. The dotted line represents the region where the TFBS sequence starts. The primers used to amplify the TFBS sequences are shown as red arrows in the schematic diagrams, with one of the primers being located within the inserted TFBS and the other one immediately outside. The grey shaded area represent a nucleosome depleted region. **d**, Scatter plots showing population-averaged nucleosome occupancy (red) and DNA methylation (black) levels within the TFBS sequence in cells with either the PE *Sox1(+35)TFBS* (left panel) or PE *Sox1(+35)TFBS+CGI* (right panel) modules inserted within the *Gata6*-TAD. The grey shaded area represent a nucleosome depleted region.

**Extended Data Fig.8.**
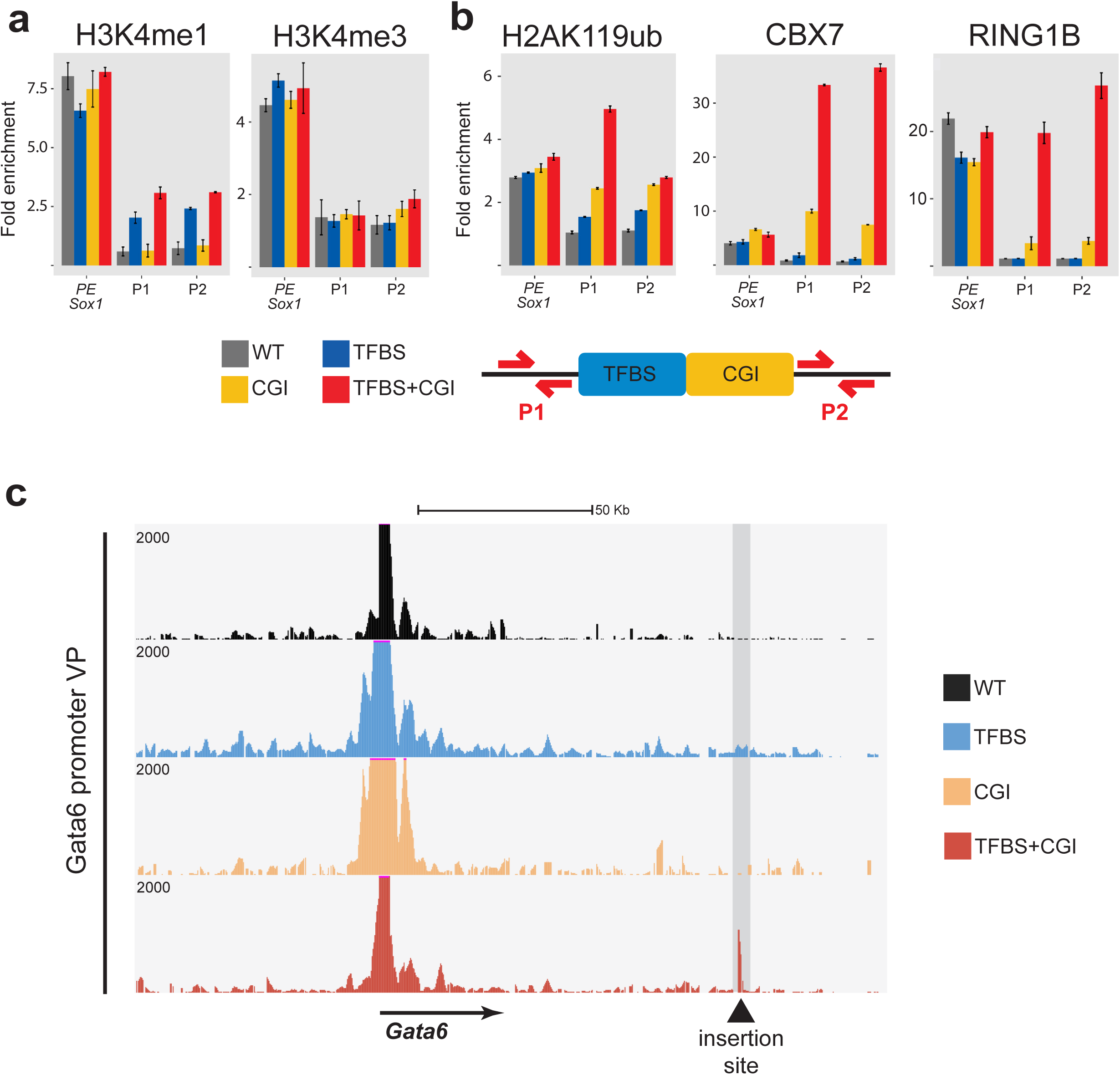
Chromatin and topological features of the PE Sox1(+35) modules engineered within the Gata6-TAD. **a-b**, H3K4me1 (a), H3K4me3 (a), H2AK119ub (b), CBX7 (b) and RING1B (b) levels at the endogenous *PE Sox1(+35)* and the *Gata6-*TAD insertion site (primer pairs P1 and P2) were measured by ChIP-qPCR in mESCs that were either WT (gray) or homozygous for the insertions of the different *PE Sox1(+35)* modules (i.e. TFBS (blue), CGI (yellow), TFBS+CGI (red)). ChIP-qPCR signals were normalized against two negative control regions (Supplementary Data 1). Error bars correspond to standard deviations from technical triplicates. The primers P1 and P2 around the *Gata6-*TAD insertion site are described in Fig 3b. c, 4C-seq experiments were performed using the *Gata6* promoter as a viewpoint in AntNPC that were either WT (black) or homozygous for the insertions of the different *PE Sox1(+35)* modules (i.e. TFBS (blue), CGI (yellow), TFBS+CGI (red)).

**Extended Data Fig.9.**
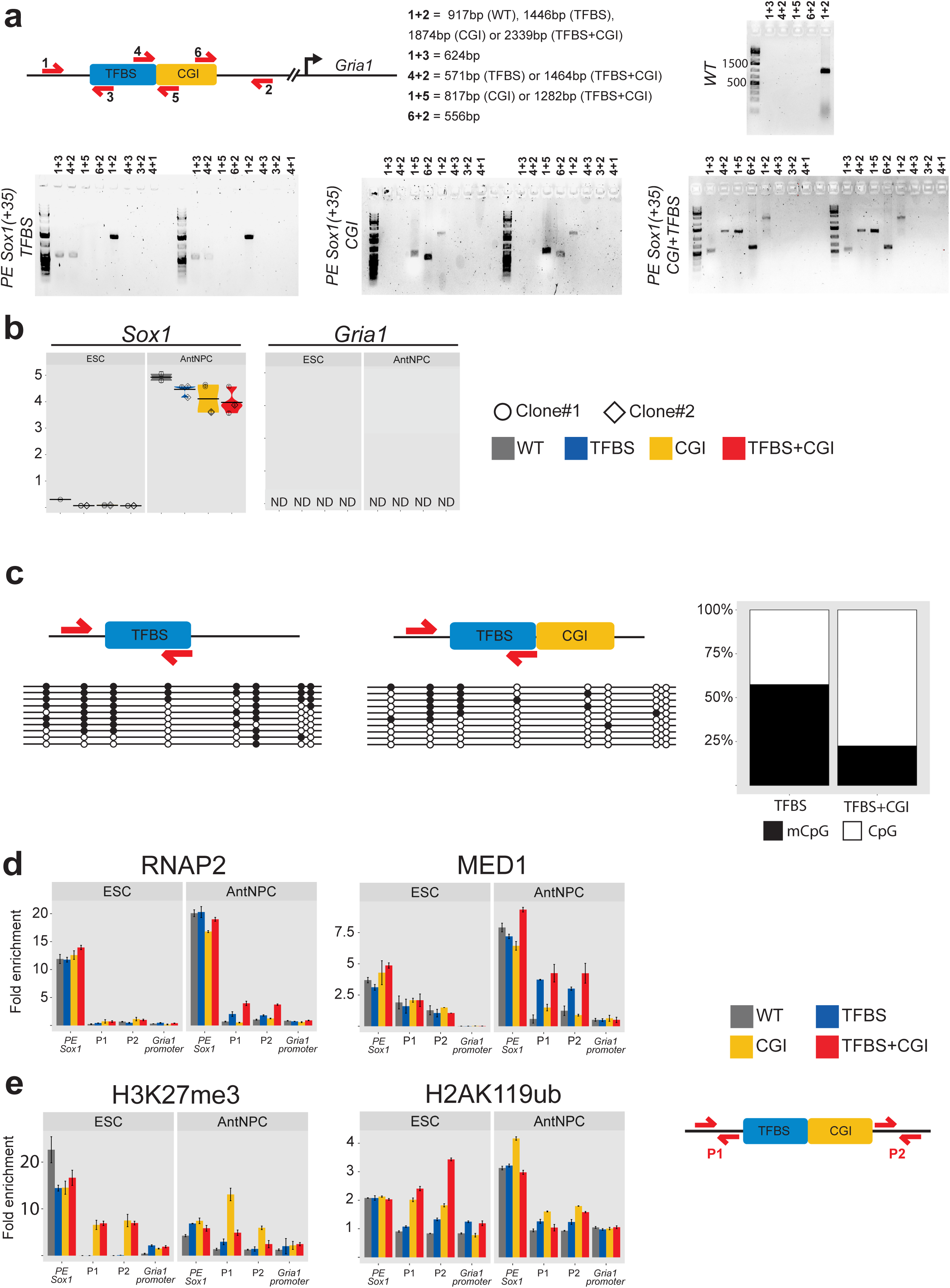
Generation and characterization of cell lines with engineered *PE Sox1(+35)* modules within the *Gria1-TAD*. a, For the identification of mESC clonal lines with the desired insertion of the different *PE Sox1(+35)* modules, primer pairs flanking the insertion borders (1+3 and 4+2; 1+5 and 6+2; or 1+3 and 6+2), amplifying potential duplications (4+3, 3+2 and 4+1; or 6+5, 5+2 and 6+1) and amplifying a large or small fragment depending on the absence or presence of the insertion (1+2), respectively, were used. The PCR results obtained for WT mESC or two mESC clonal lines with homozygous insertions of the different *PE Sox1(+35)* modules (i.e. (i) *PE Sox1(+35)TFBS*; (ii) *PE Sox1(+35)CGI*; (iii) *PE Sox1(+35)TFBS+CGI*) in the *Gria1-*TAD are shown. b, Independent biological replicate for the data presented in Fig. 4b. The expression of *Gria1* and *Sox1* was measured by RT-qPCR in mESCs (left panels) and AntNPC (right panels) that were either WT (grey) or homozygous for the insertions of the different *PE Sox1(+35)* modules (TFBS (blue), CGI (yellow), TFBS+CGI (red)). For the cells with the PE module insertions, two different clonal lines (circles and diamonds) were studied in each case. For each cell line, two technical replicates of the AntNPC differentiation were performed. The plotted expression values for each clone correspond to the average and standard deviation (error bars) from three RT-qPCR technical replicates. Expression values were normalized to two housekeeping genes (*Eef1a* and *Hprt*). c, Bisulfite sequencing analyses of ESC lines with the PE *Sox1(+35)*TFBS or *PE Sox1(+35)TFBS+CGI* modules inserted in the *Gria1-*TAD. DNA methylation levels were measured using a forward bisulfite primer upstream of the insertion site and a reverse primer inside the TFBS module (see Methods). The circles shown in the left plots correspond to individual CpG dinucleotides located within the TFBS module: unmethylated CpGs are shown in white, methylated CpGs in black and not-covered CpGs are shown in gray. 10 alleles (rows) were analyzed for each differentiation stage and cell line. The plot on the right summarizes the DNA methylation levels measured within the TFBS in the mESC lines containing the PE *Sox1(+35)TFBS* and *PE Sox1(+35)TFBS+CGI* mESC inserts within the *Gria1*-TAD. d-e, RNAP2 (d), MED1 (d), H3K27me3 (e) and H2AK119ub (e) levels at the endogenous *PE Sox1(+35)*, the *Gria1-*TAD insertion site (primer pairs P1 and P2) and the *Gria1* promoter were measured by ChIP-qPCR in mESCs (left panels) and AntNPC (right panels) that were either WT (gray) or homozygous for the insertions of the different *PE Sox1(+35)* modules (i.e. TFBS (blue), CGI (yellow), TFBS+CGI (red)). ChIP-qPCR signals were normalized against two negative control regions (Supplementary Data 1). Error bars correspond to standard deviations from technical triplicates. The location of the primers P1 and P2 around the *Gria1-*TAD insertion site is represented as red arrows in the diagram shown to the right.

**Extended Data Fig.10.**
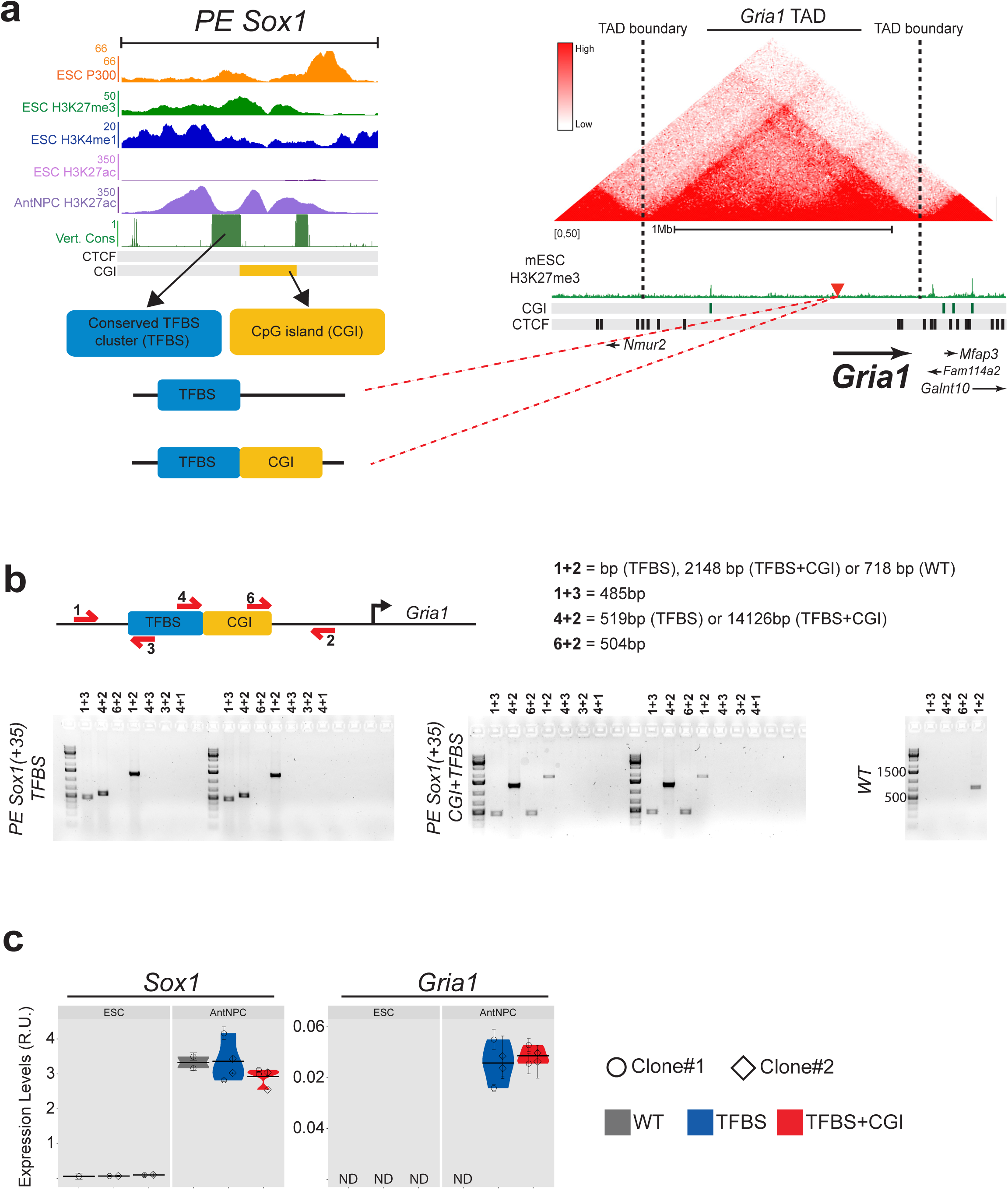
Modular engineering of the *PE Sox1(+35)* immediately upstream of the *Gria1*-TSS. a, Strategy used to insert the *PE Sox1(+35)TFBS* and *PE Sox1(+35)TFBS+CGI* modules 380 bp upstream of the *Gria1* TSS. The upper left panel shows a close-up view of the epigenomic and genetic features of the *PE Sox1(+35)*. The represented CGIs correspond to those computationally defined in the UCSC browser according to the following criteria: GC content > 50%; Length > 200 bp; CpG Observed to expected ratio > 0.6. The lower left panel shows the two combinations of *PE Sox1(+35)* modules (i.e. *PE Sox1(+35)TFBS* and *PE Sox1(+35)TFBS&CGI)* inserted 380 bp upstream of the *Gria1* TSS. The right panel shows the TAD in which *Gria1* is included (i.e. *Gria1*-TAD) according to publically available Hi-C data^33, 65^; TAD boundaries are denoted with dotted lines; H3K27me3 ChIP-seq signals in mESC are shown in green^9^; CGIs are indicated as green rectangles; CTCF binding sites^35^ are indicated as black rectangles; the yellow triangle indicates the integration site of the *PE Sox1(+35)* modules, 380 bp upstream of the *Gria1* TSS. b, For the identification of mESC clonal lines with the desired insertion of the different *PE Sox1(+35)* modules, primer pairs flanking the insertion borders (1+3 and 4+2; or 1+3 and 6+2), amplifying potential duplications (4+3, 3+2 and 4+1) and amplifying a large or small fragment depending on the absence or presence of the insertion (1+2), respectively, were used. The PCR results obtained for WT mESC or two mESC clonal lines with homozygous insertions of the different *PE Sox1(+35)* modules (i.e. (i) *PE Sox1(+35)TFBS*; (ii) *PE Sox1(+35)TFBS+CGI*) 380 bp upstream of the *Gria1* TSS are shown. c, Independent biological replicate for the data presented in Fig. 4f. The expression of *Gria1* and *Sox1* was measured by RT-qPCR in mESCs (left panels) and AntNPC (right panels) that were either WT (grey) or homozygous for the insertions of the different *PE Sox1(+35)* modules 380 bp upstream of the *Gria1* TSS (TFBS (blue); TFBS+CGI (red)). For the cells with the PE module insertions, two different clonal lines (circles and diamonds) were studied in each case. For each cell line, two technical replicates of the AntNPC differentiation were performed. The plotted expression values for each clone correspond to the average and standard deviation (error bars) from three RT-qPCR technical replicates. Expression values were normalized to two housekeeping genes (*Eef1a* and *Hprt*).

**Extended Data Fig.11.**
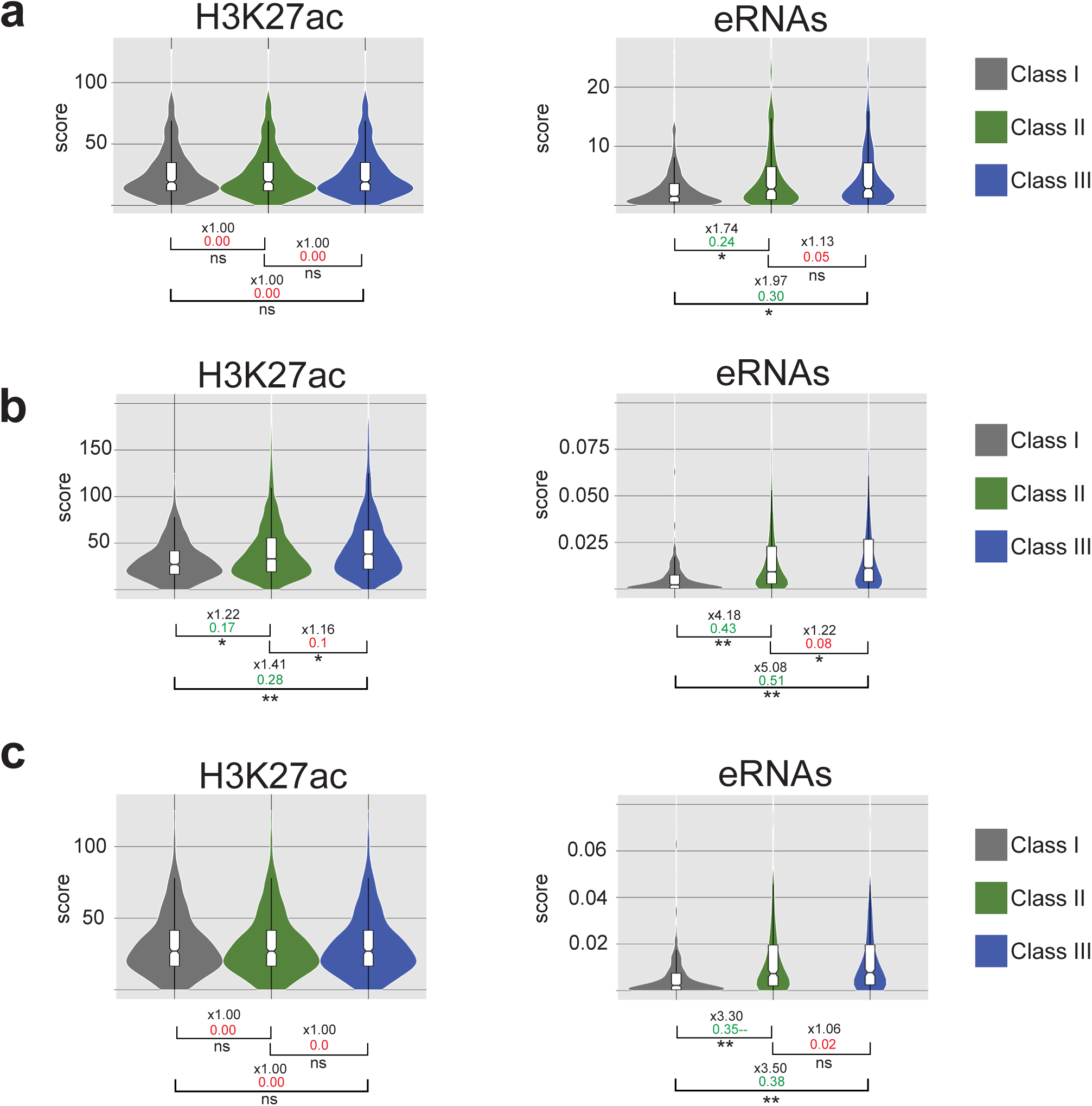
H3K27ac and eRNA levels for different classes of active enhancers in mESC. a-c, Active enhancers identified in mESC based on the presence of distal H3K27ac peaks (see Methods) were classified into three categories: *Class I* correspond to active enhancers located in TADs containing only poorly expressed genes (all genes with <0.5 FPKM); *Class II* correspond to active enhancers located in a TAD with at least one gene with expression levels >10 FPKM; *Class III* correspond to active enhancers whose closest gene in their same TAD has expression levels >10 FPKM). The violin plots show the H3K27ac (left) and eRNA (right) levels for each of these active enhancer categories in mESC. On the bottom of each plot, the asterisks indicate P-values calculated using unpaired Wilcoxon tests with Bonferroni correction for multiple testing (** = p.val < 1e^-10^; * p.val < 0.05); the numbers in black indicate the median fold-changes between the indicated groups; the coloured numbers correspond to Cliff Delta effect sizes: negligible (red) and non-negligible (green). In (a), the H3K27ac ChIP-seq data was obtained from^9^ and the PRO-seq data to measured eRNA levels was obtained from^66^. In (b-c), the H3K27ac ChIP-seq data was obtained from^69^ and the PRO-seq data to measured eRNA levels was obtained from^109^. In (a) and (c), eRNA levels for the three enhancers classes are compared after correcting for H3K27ac differences (see Methods).

**Extended Data Fig.12.**
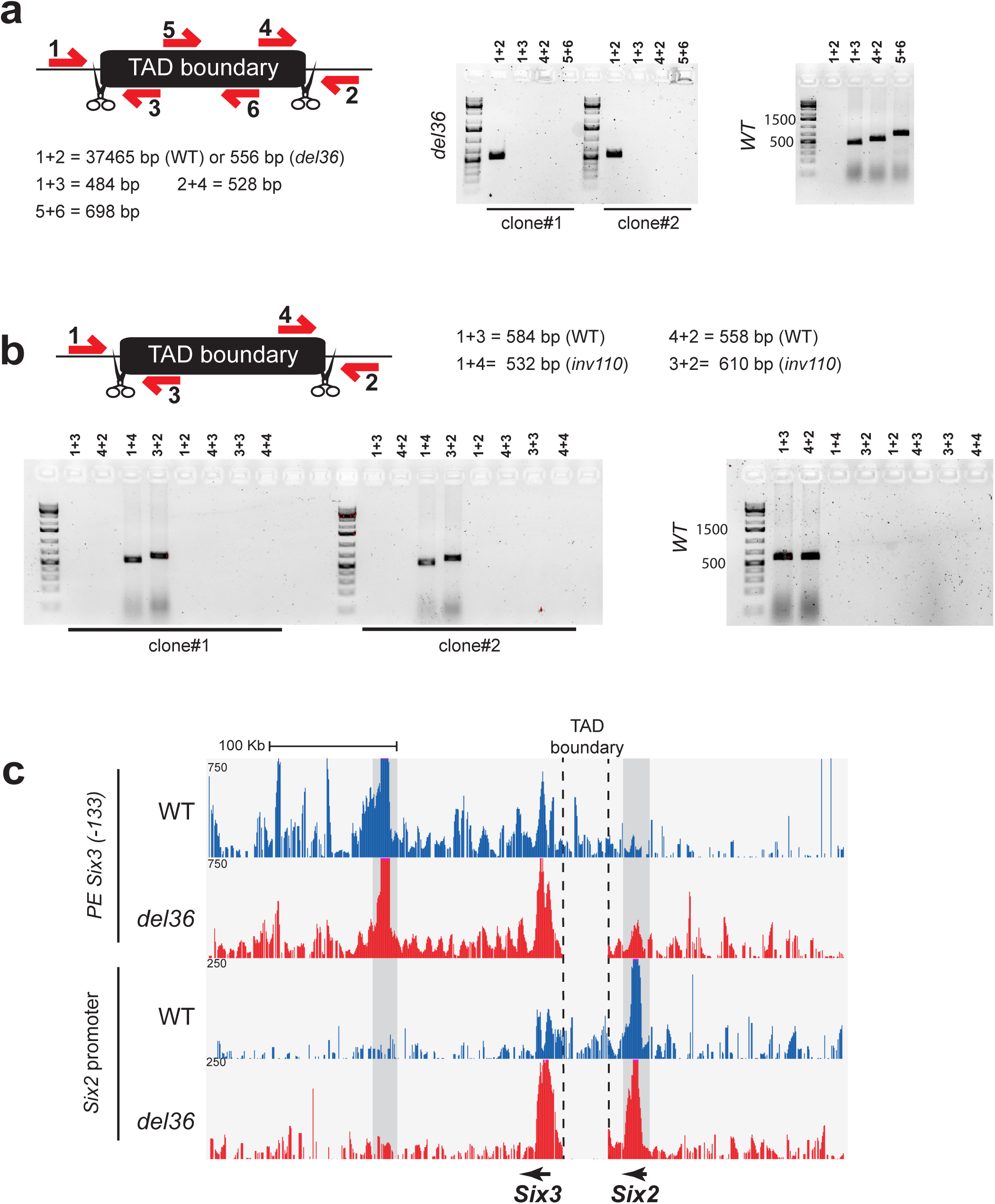
Generation of mESC lines with structural variants within the *Six3/Six2* locus. a, For the identification of mESC lines with the *Six3/Six2* TAD boundary deletion, primer pairs flanking the deleted region (1+3 and 4+2), amplifying the deleted fragment (5+6) and amplifying a large or small fragment depending on the absence or presence of the deletion (1+2), respectively, were used. The PCR results obtained for two mESC clonal lines with 36Kb homozygous deletions (*del36*) are shown. b, For the identification of mESC lines with the *Six3/Six2* inversion, primer pairs flanking the inverted region (1+3, 4+2, 1+4 and 3+2) and amplifying potential duplications (4+3, 3+3 and 4+4) were used. The PCR results obtained for two mESC clonal lines with 110 Kb homozygous inversions (*inv110*) are shown. c, 4C-seq experiments were performed using the *PE Six3(-133)* (upper panels) or the *Six2* promoter (lower panels) as viewpoints in mESCs that were either WT (blue) or homozygous for the *del36* deletion (red).

