## Supplementary Data 2 for "Orphan CpG islands boost the regulatory activity of poised enhancers and dictate the responsiveness of their target genes"

**Supplementary Data 2:** Knock-in donor templates

Legend: Homology arm TFBS module CGI module

#### *PE Sox1(+35)TFBS* in *Gata6* TAD

TCTACTGGCACATGCAGGACGACCTCAATGAGTATTTGTTGGAATTAAATGAAAGCTTGACTATGCTCAAGGAGTAGGTTGGTTCTTTGGCTGACCTCGATGAGGAA
GAGACCTTGGGTGGAAAAAGAGCAGGAGAGTGGGATCTCATAACTCACGACGACTAGTGCCAGTACCCAAAAAGCGGGGGGGTAGGAGGGAGCCGCTTGCCATTAAT
CTGGGAACAAACAGCTAACCCCGGTGACTGGTATTTTTCCTTTTCTTTTCTCACTTTTCTCAGTGTGGGGAAGGCGATTGTGAGGCGCTCTGCTGGAATTTGGCAGC
GCGGAGGCTTGGAGAGCAGCCCCATGCTGGCTCCTATTCAGCCGGCCAGTTTTCCTCGAGCTTTGGAAGTTTCACTCAGCCGTGCACTCAATGGCTTCACAAAGCTG
ATTACAAGCTTCAGCGCATTCCTGAAGGAGCCAAAAGCGACGCAGGTGCAAACGAGCCGAGGGAGCCCCTTATCCCGGTGACAGAATGGGACAAGCTGGGAAAGGCT
TGGACCACACAAATCCAAGGCTCACCAGGCAGCAGAGAGCCTGCCTTGGGAACCGGGGGTCATTATCCGCCCTATTCAGCGGGACCGGGGACCCTGGGGCCAAGGAG
GCCGGCCGGGCGCCAGTACCCAAAAAGCGGGGGGACGCGTTCCTGGAACCTCCCCTAAGCCCCTCTGGCTTCAGCTCTATTTGAGATGGATTCAGTGTCAAGGGGAA
GACAGAATTTGAGTAGAGTGAAGTCTTCAATAAGAACTGCTGCCCA

#### *PE Sox1(+35)CGI* in *Gata6* TAD

TCTACTGGCACATGCAGGACGACCTCAATGAGTATTTGTTGGAATTAAATGAAAGCTTGACTATGCTCAAGGAGTAGGTTGGTTCTTTGGCTGACCTCGATGAGGAA
GAGACCTTGGGTGGAAAAAGAGCAGGAGAGTGGGATCTCATAACTCACGACGACTAGTGCCAGTACCCAAAAAGCGGGGGGATTATCCGCCCTATTCAGCGGGACCG
GGGACCCTGGGGCCAAGGAGGCCGGCCGGGCGCAAGCGCCACCGCCAGTACGCGCCTGCAGCAGTGGGGACCCCGCTGGGCGTTGCAACCCGTCGGGATGTTGGGCG
CTGCGTTTGTTCCCACCGACAGCGGTCAGCGAAATCCATCCTGCGGGTGAAAGCCGGTGTCCAGGCTAACCGGGATACAGGAGAAGGAGGGTAGCGAAGATGCTCAG
TGCGTGGACAGTGCGTAACTTTGGTAATCGAGGCCGCACTGACGTGGTCCTGCCTCAGGCCACTCTCAGCCCCATCCTATCCCTGCTCGGGCCCGGTCCCCCTCAGC
TTGTCTTCATTTTTCGTCTCTATGAGTCATACTGAGAGCGACCTTCAGAAAGTGCGGTGGGTGGCGGTGGGTGGGTGGGTGGGGGGTTCAGGGGCGGGGAATACTGG
CTGGTAGCCCCGCAGGATCAGATCACCAGGTTGTCCAGTGCACCCTGCGGGGCAGTGGCGAGCCTGGAGAGCCGCCTCTGCCTGGGAGAATTAACGACGTCACAGCC
AGAGACGAAAGAACTACAGGTCGCGGGGACAGGCTGGCTCCCAGGGTAGAGCTTGCAAAAGGGACGAGCGTGGGACAGACTGAGGTGGGACTGTTGTAGTTCAGGGG
CCGGCCGATCCGGCTCCGTAGGCTGCCCCTATGTCCCTCCCTCGCCCCATCCAGGCCAGCACAGCACGAGTAACGCCAGCCTAGAGCCCTCCCGCCGCGTCCTCCCA
CTCCCGGCCCCACGTCTCGAAGCCACAGCCCAGCCCCACGCTTTGCCCGGCTGCACCCCGAGGTGCCCGCGTGCAGGGCGCGGGCCGGCAGGGCGCGCTTGCCGACG
CTCTTTTGTCTGCCAGTACCCAAAAAGCGGGGGGACGCGTTCCTGGAACCTCCCCTAAGCCCCTCTGGCTTCAGCTCTATTTGAGATGGATTCAGTGTCAAGGGGAA
GACAGAATTTGAGTAGAGTGAAGTCTTCAATAAGAACTGCTGCCCA

#### *PE Sox1(+35)TFBS+CGI* in *Gata6* TAD

TCTACTGGCACATGCAGGACGACCTCAATGAGTATTTGTTGGAATTAAATGAAAGCTTGACTATGCTCAAGGAGTAGGTTGGTTCTTTGGCTGACCTCGATGAGGAA
GAGACCTTGGGTGGAAAAAGAGCAGGAGAGTGGGATCTCATAACTCACGACGACTAGTGCCAGTACCCAAAAAGCGGGGGGGTAGGAGGGAGCCGCTTGCCATTAAT
CTGGGAACAAACAGCTAACCCCGGTGACTGGTATTTTTCCTTTTCTTTTCTCACTTTTCTCAGTGTGGGGAAGGCGATTGTGAGGCGCTCTGCTGGAATTTGGCAGC
GCGGAGGCTTGGAGAGCAGCCCCATGCTGGCTCCTATTCAGCCGGCCAGTTTTCCTCGAGCTTTGGAAGTTTCACTCAGCCGTGCACTCAATGGCTTCACAAAGCTG
ATTACAAGCTTCAGCGCATTCCTGAAGGAGCCAAAAGCGACGCAGGTGCAAACGAGCCGAGGGAGCCCCTTATCCCGGTGACAGAATGGGACAAGCTGGGAAAGGCT
TGGACCACACAAATCCAAGGCTCACCAGGCAGCAGAGAGCCTGCCTTGGGAACCGGGGGTCATTATCCGCCCTATTCAGCGGGACCGGGGACCCTGGGGCCAAGGAG
GCCGGCCGGGCGCAATCGATGTCATTATCCGCCCTATTCAGCGGGACCGGGGACCCTGGGGCCAAGGAGGCCGGCCGGGCGCAAGCGCCACCGCCAGTACGCGCCTG
CAGCAGTGGGGACCCCGCTGGGCGTTGCAACCCGTCGGGATGTTGGGCGCTGCGTTTGTTCCCACCGACAGCGGTCAGCGAAATCCATCCTGCGGGTGAAAGCCGGT
GTCCAGGCTAACCGGGATACAGGAGAAGGAGGGTAGCGAAGATGCTCAGTGCGTGGACAGTGCGTAACTTTGGTAATCGAGGCCGCACTGACGTGGTCCTGCCTCAG
GCCACTCTCAGCCCCATCCTATCCCTGCTCGGGCCCGGTCCCCCTCAGCTTGTCTTCATTTTTCGTCTCTATGAGTCATACTGAGAGCGACCTTCAGAAAGTGCGGT
GGGTGGCGGTGGGTGGGTGGGTGGGGGGTTCAGGGGCGGGGAATACTGGCTGGTAGCCCCGCAGGATCAGATCACCAGGTTGTCCAGTGCACCCTGCGGGGCAGTGG
CGAGCCTGGAGAGCCGCCTCTGCCTGGGAGAATTAACGACGTCACAGCCAGAGACGAAAGAACTACAGGTCGCGGGGACAGGCTGGCTCCCAGGGTAGAGCTTGCAA
AAGGGACGAGCGTGGGACAGACTGAGGTGGGACTGTTGTAGTTCAGGGGCCGGCCGATCCGGCTCCGTAGGCTGCCCCTATGTCCCTCCCTCGCCCCATCCAGGCCA
GCACAGCACGAGTAACGCCAGCCTAGAGCCCTCCCGCCGCGTCCTCCCACTCCCGGCCCCACGTCTCGAAGCCACAGCCCAGCCCCACGCTTTGCCCGGCTGCACCC
CGAGGTGCCCGCGTGCAGGGCGCGGGCCGGCAGGGCGCGCTTGCCGACGCTCTTTTGTCTGCCAGTACCCAAAAAGCGGGGGGACGCGTTCCTGGAACCTCCCCTAA
GCCCCTCTGGCTTCAGCTCTATTTGAGATGGATTCAGTGTCAAGGGGAAGACAGAATTTGAGTAGAGTGAAGTCTTCAATAAGAACTGCTGCCCA

#### *PE Sox1(+35)TFBS+aCGI* in *Gata6* TAD

TCTACTGGCACATGCAGGACGACCTCAATGAGTATTTGTTGGAATTAAATGAAAGCTTGACTATGCTCAAGGAGTAGGTTGGTTCTTTGGCTGACCTCGATGAGGAA
GAGACCTTGGGTGGAAAAAGAGCAGGAGAGTGGGATCTCATAACTCACGACGACTAGTGCCAGTACCCAAAAAGCGGGGGGGTAGGAGGGAGCCGCTTGCCATTAAT
CTGGGAACAAACAGCTAACCCCGGTGACTGGTATTTTTCCTTTTCTTTTCTCACTTTTCTCAGTGTGGGGAAGGCGATTGTGAGGCGCTCTGCTGGAATTTGGCAGC
GCGGAGGCTTGGAGAGCAGCCCCATGCTGGCTCCTATTCAGCCGGCCAGTTTTCCTCGAGCTTTGGAAGTTTCACTCAGCCGTGCACTCAATGGCTTCACAAAGCTG
ATTACAAGCTTCAGCGCATTCCTGAAGGAGCCAAAAGCGACGCAGGTGCAAACGAGCCGAGGGAGCCCCTTATCCCGGTGACAGAATGGGACAAGCTGGGAAAGGCT
TGGACCACACAAATCCAAGGCTCACCAGGCAGCAGAGAGCCTGCCTTGGGAACCGGGGGTCATTATCCGCCCTATTCAGCGGGACCGGGGACCCTGGGGCCAAGGAG
GCCGGCCGGGCGCAATCGATGTCCCCAGGTCAGGAGCCGCGGCTCCTTCCTACCGTCGTGCCCAGGAGGTAGGGTTGGACACCCTCCGGTAGGTGGCGTCAGACCCC
ACACGCGAGCGTCGTCACGCGCCGACCCTACTGTTGGCCCGTGGAGGGTGGAGTGGGGGCAGCGCACCCAGACCTGCGGGTCACCACCGGGCTCGTGGGACCAGAGG
TAGGCCTGGGAGCTCACCTCGAGGGAACCCGGCGCCACCCGCTCCCCTTAGCCGTGGGACCACCTCCGACCCCAGGCCTGGGTTGCCACTACCGGCGGTCCTAAGGG
GAGGTGGGCACGGCTCCACGTGGCGCGAGGGTCCTGTGGGCAGGTCCCTCCCTGGCCAGCAACGGGGGCTAGGCTAGGAGCCCGGCACTCTGGCCAGCCTTAGGCG
ACCCCCACCTAGGCCTACCTGTTCGGGCACGACCCCTGGCTGCGCTGGCGACCGCCACCACTGCCAGCGTCCTCGAGAGTGCCGTCCAGGCTGGGTCCGGGCTCCGA
GGCCTGCTGTGGCCTCGCAGACCACCGCTGGGCCAAACACCCGAGGCCGGGTTGGCCCCTTTCGGAGCCCGCGACCCAGCGGCAAGGTCTGGAAGCCAGCCTGGCCC
AAGGACGGACGGGCCACCGACCGCGTCGGTTCCCCCGAGCGCGGCAGGACTGGCTACAGGGCCCCTGCCGACGGCTAGGCTGGACGTTCGGAGGTGTCCTAGACTTC

CCATCGGGGCAGAGACCCGGAGTCGTGCCACCCGGGTTTCAGGAGTGACGGTGGGACAACGCGTTCCTGGAACCTCCCCTAAGCCCCTCTGGCTTCAGCTCTATTTG
AGATGGATTCAGTGTCAAGGGGAAGACAGAATTTGAGTAGAGTGAAGTCTTCAATAAGAACTGCTGCCCA

### *PE Wnt8b(+21)TFBS* in *Gata6* TAD

TCTACTGGCACATGCAGGACGACCTCAATGAGTATTTGTTGGAATTAAATGAAAGCTTGACTATGCTCAAGGAGTAGGTTGGTTCTTTGGCTGACCTCGATGAGGAA
GAGACCTTGGGTGGAAAAAGAGCAGGAGAGTGGGATCTCATAACTCACGACGACTAGTGATCCTCCCTCCTCTTGTGGGCTCTTAAGGAGCCCATCTGTCTCAGTCT
GAGGACAGAAGGAGAGGGGGGGTCCTGAGAAGTGACAAGAACTCTGGGTAGGGTGAGCAAAACACTGTCTCCTTCCTGCCCCTGGCTGGTGAGCCTACACATAAGCA
GGGCTTTCTTTTCTTTAAAAGAATCCCACCCTCACTTGGCCTCAGGAGCAGGTCAGTGCGGAGAGAGAAAGGCTCCTCCTTGGCACGCTGGTAGCGCTGGTGGTGGT
GGGAGGGAGGGCAGGCCAGGGCACTCGCCTATTGTGGGTCTGGATTGGATGCTTCCACTCATTCGTTTCCTTTTAATTAACCCACATTCCTGGAGGAGAAATATTTA
TGTTGGCTCCCATAGAGGCAACCCTCAGACTGGAAAGAAGGGGGAAGAAAGCCTGTCTTAGCAAAACAATATCTCTAAAGAGCTCAAGAAGAGAAAATAGGGGAACA
AAATGCCATTAAAACAGCCTTCTACCCCCGCCGGAGTCAATGAAAACTTCATTTCACATGTTAATCCGCCACTCCGCCCCTTCTCTTCTCACAACACCTTTTCGGAA
AGTGTTTGGGAAAAGTCTTGGCAGTCAGTACACAGCTCCCTGCAGCTCAGACTTCTCCTTGTGTTAGTTCAGGGCCCCGAGTGGGGCTCTCGCTCCAGCAGCGGCTT
TTTCTCTCCCCCTCCTTTAATTTAAACAAAGACTTCAGAGTTCATCAACCTCCCACTGAAATTCACAGCTGCTGAACAAACTAATCCCCTCTCCCCGCCTTTCTTCC
CTTCAATGGGCTCTTGGCTTCAAAGACACTTTGGGAAAGGACTTTGCTGGGGCTCACACTGCTTCATCAATAGAAGTTAGCACAAGTATTATCTGAAGAGCAAGTGG
CTTTACAAACAAATACTGCTTAACAATCCCTCCCTCCCAAACACACACATCTAGCCTGCAGACATGATGGATGGACAACAGGATTAGGCCTGGTACCCCCTGGCTTC
ACACCCAGGCCCTTTGTGCAGCTGAACTCAGCCCTGAAGTGGTGTGGGGGAAACCCTGCAGCTCACTGTCAGGCACAGATGGAGACAGACAACGCGTTCCTGGAACC
TCCCCTAAGCCCCTCTGGCTTCAGCTCTATTTGAGATGGATTCAGTGTCAAGGGGAAGACAGAATTTGAGTAGAGTGAAGTCTTCAATAAGAACTGCTGCCCA

### *PE Wnt8b(+21)CGI* in *Gata6* TAD

TCTACTGGCACATGCAGGACGACCTCAATGAGTATTTGTTGGAATTAAATGAAAGCTTGACTATGCTCAAGGAGTAGGTTGGTTCTTTGGCTGACCTCGATGAGGAA
GAGACCTTGGGTGGAAAAAGAGCAGGAGAGTGGGATCTCATAACTCACGACGACTAGTCCAATCGTCTGGCCACTCACTAGCTAGGCGCCTGGCAACTGCTCCAGCT
CGGGTGGCTTACGTGCTGCTCTCTCCCCCAGGCGGTCAAGGGCACCATGAAACGCACGTGTAAGTGCCACGGTGTGTCCGGCAGCTGCACCACGCAGACCTGCTGGT
TGCAACTGCCAGAGTTCCGGGAGGTAGGCGCGCACTTGAAGGAGAAGTATCATGCGGCGCTCAAGGTGGACCTGCTGCAAGGCGCGGGCAACAGCGCGGCGGGCCGC
GGAGCCATCGCCGACACCTTCCGCTCCATCTCCACCCGCGAGCTGGTGCATCTGGAGGACTCCCCAGACTACTGCCTGGAGAACAAGACCCTGGGGCTGCTGGGCAC
CGAGGGCCGAGAGTGTCTGCGGCGCGGGCGCGCCCTGGGTCGCTGGGAGCGCCGCAGTTGTCGCCGGCTGTGCGGGGACTGCGGGCTAGCGGTGGAGGAGCGCCGCG
CCGAGACAGTGTCCAGCTGCAACTGCAAGTTTCACTGGTGCTGCGCGGTCCGCTGCGAGCAGTGCCGCCGGCGGGTCACCAAGTACTTCTGCAGCCGCGCAGAGCGG
CCGCCCAGAGGCGCTGCGCACAAACCGGGAAAGAACTCCTAAGGGTATCTATCCCTCCCGCCTCCACCCCTGTTCGTCCTCGGCTTCCTTTAGAGACCCGGATCACG
CGTTCCTGGAACCTCCCCTAAGCCCCTCTGGCTTCAGCTCTATTTGAGATGGATTCAGTGTCAAGGGGAAGACAGAATTTGAGTAGAGTGAAGTCTTCAATAAGAAC
TGCTGCCCA

### *PE Wnt8b(+21)TFBS+CGI* in *Gata6* TAD

TCTACTGGCACATGCAGGACGACCTCAATGAGTATTTGTTGGAATTAAATGAAAGCTTGACTATGCTCAAGGAGTAGGTTGGTTCTTTGGCTGACCTCGATGAGGAA
GAGACCTTGGGTGGAAAAAGAGCAGGAGAGTGGGATCTCATAACTCACGACGACTAGTCCAATCGTCTGGCCACTCACTAGCTAGGCGCCTGGCAACTGCTCCAGCT
CGGGTGGCTTACGTGCTGCTCTCTCCCCCAGGCGGTCAAGGGCACCATGAAACGCACGTGTAAGTGCCACGGTGTGTCCGGCAGCTGCACCACGCAGACCTGCTGGT
TGCAACTGCCAGAGTTCCGGGAGGTAGGCGCGCACTTGAAGGAGAAGTATCATGCGGCGCTCAAGGTGGACCTGCTGCAAGGCGCGGGCAACAGCGCGGCGGGCCGC
GGAGCCATCGCCGACACCTTCCGCTCCATCTCCACCCGCGAGCTGGTGCATCTGGAGGACTCCCCAGACTACTGCCTGGAGAACAAGACCCTGGGGCTGCTGGGCAC
CGAGGGCCGAGAGTGTCTGCGGCGCGGGCGCGCCCTGGGTCGCTGGGAGCGCCGCAGTTGTCGCCGGCTGTGCGGGGACTGCGGGCTAGCGGTGGAGGAGCGCCGCG
CCGAGACAGTGTCCAGCTGCAACTGCAAGTTTCACTGGTGCTGCGCGGTCCGCTGCGAGCAGTGCCGCCGGCGGGTCACCAAGTACTTCTGCAGCCGCGCAGAGCGG
CCGCCCAGAGGCGCTGCGCACAAACCGGGAAAGAACTCCTAAGGGTATCTATCCCTCCCGCCTCCACCCCTGTTCGTCCTCGGCTTCCTTTAGAGACCCGGATCCTC
CCTCCTCTTGTGGGCTCTTAAGGAGCCCATCTGTCTCAGTCTGAGGACAGAAGGAGAGGGGGGGTCCTGAGAAGTGACAAGAACTCTGGGTAGGGTGAGCAAAACAC
TGTCTCCTTCCTGCCCCTGGCTGGTGAGCCTACACATAAGCAGGGCTTTCTTTTCTTTAAAAGAATCCCACCCTCACTTGGCCTCAGGAGCAGGTCAGTGCGGAGAG
AGAAAGGCTCCTCCTTGGCACGCTGGTAGCGCTGGTGGTGGTGGGAGGGAGGGCAGGCCAGGGCACTCGCCTATTGTGGGTCTGGATTGGATGCTTCCACTCATTCG
TTTCCTTTTAATTAACCCACATTCCTGGAGGAGAAATATTTATGTTGGCTCCCATAGAGGCAACCCTCAGACTGGAAAGAAGGGGGAAGAAAGCCTGTCTTAGCAAA
ACAATATCTCTAAAGAGCTCAAGAAGAGAAAATAGGGGAACAAAATGCCATTAAAACAGCCTTCTACCCCCGCCGGAGTCAATGAAAACTTCATTTCACATGTTAAT
CCGCCACTCCGCCCCTTCTCTTCTCACAACACCTTTTCGGAAAGTGTTTGGGAAAAGTCTTGGCAGTCAGTACACAGCTCCCTGCAGCTCAGACTTCTCCTTGTGTT
AGTTCAGGGCCCCGAGTGGGGCTCTCGCTCCAGCAGCGGCTTTTTCTCTCCCCCTCCTTTAATTTAAACAAAGACTTCAGAGTTCATCAACCTCCCACTGAAATTCA
CAGCTGCTGAACAAACTAATCCCCTCTCCCCGCCTTTCTTCCCTTCAATGGGCTCTTGGCTTCAAAGACACTTTGGGAAAGGACTTTGCTGGGGCTCACACTGCTTC
ATCAATAGAAGTTAGCACAAGTATTATCTGAAGAGCAAGTGGCTTTACAAACAAATACTGCTTAACAATCCCTCCCTCCCAAACACACACATCTAGCCTGCAGACAT
GATGGATGGACAACAGGATTAGGCCTGGTACCCCCTGGCTTCACACCCAGGCCCTTTGTGCAGCTGAACTCAGCCCTGAAGTGGTGTGGGGGAAACCCTGCAGCTCA
CTGTCAGGCACAGATGGAGACAGACAACGCGTTCCTGGAACCTCCCCTAAGCCCCTCTGGCTTCAGCTCTATTTGAGATGGATTCAGTGTCAAGGGGAAGACAGAAT
TTGAGTAGAGTGAAGTCTTCAATAAGAACTGCTGCCCA

### *PE Sox1(+35)TFBS* in *Foxa2* TAD

AGGGCATCAGCTCAGGATGATTCAAAGATTAAAGAGGTGTGAGAACAAATGCCTTTCTCTTAACCAAGCTGTCCTGTCCTGTCCTGTCCTCAGGGTTTGGGCCCAAA
CCATTCTGTGGCCTTGTATGTGGTCACAGGCTAGACCTGCAGAAGAGAATTTTAAGGAACAGATGAAAATGACAGTGAGGGCTCTATGATAAGATCACTAGTGCCAG
TACCCAAAAAGCGGGGGGTAGGAGGGAGCCGCTTGCCATTAATCTGGGAACAAACAGCTAACCCCGGTGACTGGTATTTTTCCTTTTCTTTTCTCACTTTTCTCAGT
GTGGGGAAGGCGATTGTGAGGCGCTCTGCTGGAATTTGGCAGCGCGGAGGCTTGGAGAGCAGCCCCATGCTGGCTCCTATTCAGCCGGCCAGTTTTCCTCGAGCTTT
GGAAGTTTCACTCAGCCGTGCACTCAATGGCTTCACAAAGCTGATTACAAGCTTCAGCGCATTCCTGAAGGAGCCAAAAGCGACGCAGGTGCAAACGAGCCGAGGGA
GCCCCTTATCCCGGTGACAGAATGGGACAAGCTGGGAAAGGCTTGGACCACACAAATCCAAGGCTCACCAGGCAGCAGAGAGCCTGCCTTGGGAACCGGGGGTCATT
ATCCGCCCTATTCAGCGGGACCGGGGACCCTGGGGCCAAGGAGGCCGGCCGGGCGCCAGTACCCAAAAAGCGGGGGGACGCGTCTGAGGCCCCTGGATTAGCTCCCA
GCAATAAATTTCATAACTTGCTACAGTATTCACGCTTACTACTTGAAAAAAACACTGCAAGGAAATTTATTGAGAATGAGGAGAAAGGTGGGAGAATGACTAAGATA
GGAGAATTGCTTTGGGTTTTTTTGTTTATAAACAATGACCATAAAAGCAAGGACCAGGGTAGGGTGTGGAGTGAAATAATGAGCCATTTGCCTAATGATTATT

### *PE Sox1(+35)CGI* in *Foxa2* TAD

AGGGCATCAGCTCAGGATGATTCAAAGATTAAAGAGGTGTGAGAACAAATGCCTTTCTCTTAACCAAGCTGTCCTGTCCTGTCCTGTCCTCAGGGTTTGGGCCCAAA
CCATTCTGTGGCCTTGTATGTGGTCACAGGCTAGACCTGCAGAAGAGAATTTTAAGGAACAGATGAAAATGACAGTGAGGGCTCTATGATAAGATCACTAGTGCCAG
TACCCAAAAAGCGGGGGATTATCCGCCCTATTCAGCGGGACCGGGGACCCTGGGGCCAAGGAGGCCGGCCGGGCGCAAGCGCCACCGCCAGTACGCGCCTGCAGCAG
TGGGGACCCCGCTGGGCGTTGCAACCCGTCGGGATGTTGGGCGCTGCGTTTGTTCCCACCGACAGCGGTCAGCGAAATCCATCCTGCGGGTGAAAGCCGGTGTCCAG
GCTAACCGGGATACAGGAGAAGGAGGGTAGCGAAGATGCTCAGTGCGTGGACAGTGCGTAACTTTGGTAATCGAGGCCGCACTGACGTGGTCCTGCCTCAGGCCACT
CTCAGCCCCATCCTATCCCTGCTCGGGCCCGGTCCCCCTCAGCTTGTCTTCATTTTTCGTCTCTATGAGTCATACTGAGAGCGACCTTCAGAAAGTGCGGTGGGTGG
CGGTGGGTGGGTGGGTGGGGGGTTCAGGGGCGGGGAATACTGGCTGGTAGCCCCGCAGGATCAGATCACCAGGTTGTCCAGTGCACCCTGCGGGGCAGTGGCGAGCC
TGGAGAGCCGCCTCTGCCTGGGAGAATTAACGACGTCACAGCCAGAGACGAAAGAACTACAGGTCGCGGGGACAGGCTGGCTCCCAGGGTAGAGCTTGCAAAAGGGA
CGAGCGTGGGACAGACTGAGGTGGGACTGTTGTAGTTCAGGGGCCGGCCGATCCGGCTCCGTAGGCTGCCCCTATGTCCCTCCCTCGCCCCATCCAGGCCAGCACAG
CACGAGTAACGCCAGCCTAGAGCCCTCCCGCCGCGTCCTCCCACTCCCGGCCCCACGTCTCGAAGCCACAGCCCAGCCCCACGCTTTGCCCGGCTGCACCCCGAGGT
GCCCGCGTGCAGGGCGCGGGCCGGCAGGGCGCGCTTGCCGACGCTCTTTTGTCTGCCAGTACCCAAAAAGCGGGGGGACGCGTCTGAGGCCCCTGGATTAGCTCCCA
GCAATAAATTTCATAACTTGCTACAGTATTCACGCTTACTACTTGAAAAAAACACTGCAAGGAAATTTATTGAGAATGAGGAGAAAGGTGGAGAATGACTAAGATA
GGAGAATTGCTTTGGGTTTTTTTGTTTATAAACAATGACCATAAAAGCAAGGACCAGGGTAGGGTGTGGAGTGAAATAATGAGCCATTTGCCTAATGATTATT

### *PE Sox1(+35)TFBS+CGI* in *Foxa2* TAD

AGGGCATCAGCTCAGGATGATTCAAAGATTAAAGAGGTGTGAGAACAAATGCCTTTCTCTTAACCAAGCTGTCCTGTCCTGTCCTGTCCTCAGGGTTTGGGCCCAAA
CCATTCTGTGGCCTTGTATGTGGTCACAGGCTAGACCTGCAGAAGAGAATTTTAAGGAACAGATGAAAATGACAGTGAGGGCTCTATGATAAGATCACTAGTGCCAG
TACCCAAAAAGCGGGGGGTAGGAGGGAGCCGCTTGCCATTAATCTGGGAACAAACAGCTAACCCCGGTGACTGGTATTTTTCCTTTTCTTTTCTCACTTTTCTCAGT
GTGGGGAAGGCGATTGTGAGGCGCTCTGCTGGAATTTGGCAGCGCGGAGGCTTGGAGAGCAGCCCCATGCTGGCTCCTATTCAGCCGGCCAGTTTTCCTCGAGCTTT
GGAAGTTTCACTCAGCCGTGCACTCAATGGCTTCACAAAGCTGATTACAAGCTTCAGCGCATTCCTGAAGGAGCCAAAAGCGACGCAGGTGCAAACGAGCCGAGGGA
GCCCCTTATCCCGGTGACAGAATGGGACAAGCTGGGAAAGGCTTGGACCACACAAATCCAAGGCTCACCAGGCAGCAGAGAGCCTGCCTTGGGAACCGGGGGTCATT
ATCCGCCCTATTCAGCGGGACCGGGGACCCTGGGGCCAAGGAGGCCGGCCGGGCGCAATCGATGTCATTATCCGCCCTATTCAGCGGGACCGGGGACCCTGGGGCCA
AGGAGGCCGGCCGGGCGCAAGCGCCACCGCCAGTACGCGCCTGCAGCAGTGGGGACCCCGCTGGGCGTTGCAACCCGTCGGGATGTTGGGCGCTGCGTTTGTTCCCA
CCGACAGCGGTCAGCGAAATCCATCCTGCGGGTGAAAGCCGGTGTCCAGGCTAACCGGGATACAGGAGAAGGAGGGTAGCGAAGATGCTCAGTGCGTGGACAGTGCG
TAACTTTGGTAATCGAGGCCGCACTGACGTGGTCCTGCCTCAGGCCACTCTCAGCCCCATCCTATCCCTGCTCGGGCCCGGTCCCCCTCAGCTTGTCTTCATTTTTC
GTCTCTATGAGTCATACTGAGAGCGACCTTCAGAAAGTGCGGTGGGTGGCGGTGGGTGGGTGGGTGGGGGGTTCAGGGGCGGGGAATACTGGCTGGTAGCCCCGCAG
GATCAGATCACCAGGTTGTCCAGTGCACCCTGCGGGGCAGTGGCGAGCCTGGAGAGCCGCCTCTGCCTGGGAGAATTAACGACGTCACAGCCAGAGACGAAAGAACT
ACAGGTCGCGGGGACAGGCTGGCTCCCAGGGTAGAGCTTGCAAAAGGGACGAGCGTGGGACAGACTGAGGTGGGACTGTTGTAGTTCAGGGGCCGGCCGATCCGGCT
CCGTAGGCTGCCCCTATGTCCCTCCCTCGCCCCATCCAGGCCAGCACAGCACGAGTAACGCCAGCCTAGAGCCCTCCCGCCGCGTCCTCCCACTCCCGGCCCCACGT
CTCGAAGCCACAGCCCAGCCCCACGCTTTGCCCGGCTGCACCCCGAGGTGCCCGCGTGCAGGGCGCGGGCCGGCAGGGCGCGCTTGCCGACGCTCTTTTGTCTGCCA
GTACCCAAAAAGCGGGGGGACGCGTCTGAGGCCCCTGGATTAGCTCCCAGCAATAAATTTCATAACTTGCTACAGTATTCACGCTTACTACTTGAAAAAAACACTGC
AAGGAAATTTATTGAGAATGAGGAGAAAGGTGGAGAATGACTAAGATAGGAGAATTGCTTTGGGTTTTTTTGTTTATAAACAATGACCATAAAAGCAAGGACCAGG
GTAGGGTGTGGAGTGAAATAATGAGCCATTTGCCTAATGATTATT

### *PE Sox1(+35)TFBS* in *Gria1* TAD

AAGTAAAGTGATTTCTTCAAAAACAAGAAAAGGAAACATTCACTTGAGAGCATGAGCAGTGCCTCTGGGTATAGGGTAGCTCCTTCTAGGACACTGAGATATTGAAA
TCGCCCCCACACTAGTGCCAGTACCCAAAAAGCGGGGGGGTAGGAGGGAGCCGCTTGCCATTAATCTGGGAACAAACAGCTAACCCCGGTGACTGGTATTTTTCCTT
TTCTTTTCTCACTTTTCTCAGTGTGGGGAAGGCGATTGTGAGGCGCTCTGCTGGAATTTGGCAGCGCGGAGGCTTGGAGAGCAGCCCCATGCTGGCTCCTATTCAGC
CGGCCAGTTTTCCTCGAGCTTTGGAAGTTTCACTCAGCCGTGCACTCAATGGCTTCACAAAGCTGATTACAAGCTTCAGCGCATTCCTGAAGGAGCCAAAAGCGACG
CAGGTGCAAACGAGCCGAGGGAGCCCCTTATCCCGGTGACAGAATGGGACAAGCTGGGAAAGGCTTGGACCACACAAATCCAAGGCTCACCAGGCAGCAGAGAGCCT
GCCTTGGGAACCGGGGGTCATTATCCGCCCTATTCAGCGGGACCGGGGACCCTGGGGCCAAGGAGGCCGGCCGGGCGCCAGTACCCAAAAAGCGGGGGGACGCGTGG
CATCAGAATGTGGCAGATTGCGGAATATAGAAAAAAGAGGGTTACAGTTTTCTATGGGAATGAGAGGTGCCAGGACTTGAAAGAAAATAAAACCACATACTCCCTCT
GCTATATTAACTGAGAGTCCTTAAAACAAATAACTTCACGTTTA

### *PE Sox1(+35)CGI* in *Gria1* TAD

AAGTAAAGTGATTTCTTCAAAAACAAGAAAAGGAAACATTCACTTGAGAGCATGAGCAGTGCCTCTGGGTATAGGGTAGCTCCTTCTAGGACACTGAGATATTGAAA
TCGCCCCCACACTAGTGCCAGTACCCAAAAAGCGGGGGGATTATCCGCCCTATTCAGCGGGACCGGGGACCCTGGGGCCAAGGAGGCCGGCCGGGCGCAAGCGCCAC
CGCCAGTACGCGCCTGCAGCAGTGGGGACCCCGCTGGGCGTTGCAACCCGTCGGGATGTTGGGCGCTGCGTTTGTTCCCACCGACAGCGGTCAGCGAAATCCATCCT
GCGGGTGAAAGCCGGTGTCCAGGCTAACCGGGATACAGGAGAAGGAGGGTAGCGAAGATGCTCAGTGCGTGGACAGTGCGTAACTTTGGTAATCGAGGCCGCACTGA
CGTGGTCCTGCCTCAGGCCACTCTCAGCCCCATCCTATCCCTGCTCGGGCCCGGTCCCCCTCAGCTTGTCTTCATTTTTCGTCTCTATGAGTCATACTGAGAGCGAC
CTTCAGAAAGTGCGGTGGGTGGCGGTGGGTGGGTGGGTGGGGGGTTCAGGGGCGGGGAATACTGGCTGGTAGCCCCGCAGGATCAGATCACCAGGTTGTCCAGTGCA
CCCTGCGGGGCAGTGGCGAGCCTGGAGAGCCGCCTCTGCCTGGGAGAATTAACGACGTCACAGCCAGAGACGAAAGAACTACAGGTCGCGGGGACAGGCTGGCTCCC
AGGGTAGAGCTTGCAAAAGGGACGAGCGTGGGACAGACTGAGGTGGGACTGTTGTAGTTCAGGGGCCGGCCGATCCGGCTCCGTAGGCTGCCCCTATGTCCCTCCCT
CGCCCCATCCAGGCCAGCACAGCACGAGTAACGCCAGCCTAGAGCCCTCCCGCCGCGTCCTCCCACTCCCGGCCCCACGTCTCGAAGCCACAGCCCAGCCCCACGCT
TTGCCCGGCTGCACCCCGAGGTGCCCGCGTGCAGGGCGCGGGCCGGCAGGGCGCGCTTGCCGACGCTCTTTTGTCTGCCAGTACCCAAAAAGCGGGGGGACGCGTGG
CATCAGAATGTGGCAGATTGCGGAATATAGAAAAAAGAGGGTTACAGTTTTCTATGGGAATGAGAGGTGCCAGGACTTGAAAGAAAATAAAACCACATACTCCCTCT
GCTATATTAACTGAGAGTCCTTAAAACAAATAACTTCACGTTTA

### *PE Sox1(+35)TFBS+CGI* in Gria1 TAD

AAGTAAAGTGATTTCTTCAAAAACAAGAAAAGGAAACATTCACTTGAGAGCATGAGCAGTGCCTCTGGGTATAGGGTAGCTCCTTCTAGGACACTGAGATATTGAAA
TCGCCCCCACACTAGTGCCAGTACCCAAAAAGCGGGGGGGTAGGAGGGAGCCGCTTGCCATTAATCTGGGAACAAACAGCTAACCCCGGTGACTGGTATTTTTCCTT
TTCTTTTCTCACTTTTCTCAGTGTGGGGAAGGCGATTGTGAGGCGCTCTGCTGGAATTTGGCAGCGCGGAGGCTTGGAGAGCAGCCCCATGCTGGCTCCTATTCAGC
CGGCCAGTTTTCCTCGAGCTTTGGAAGTTTCACTCAGCCGTGCACTCAATGGCTTCACAAAGCTGATTACAAGCTTCAGCGCATTCCTGAAGGAGCCAAAAGCGACG

CAGGTGCAAACGAGCCGAGGGAGCCCCTTATCCCGGTGACAGAATGGGACAAGCTGGGAAAGGCTTGGACCACACAAATCCAAGGCTCACCAGGCAGCAGAGAGCCT
GCCTTGGGAACCGGGGGTCATTATCCGCCCTATTCAGCGGGACCGGGGACCCTGGGGCCAAGGAGGCCGGCCGGGCGCAATCGATGTCATTATCCGCCCTATTCAGC
GGGACCGGGGACCCTGGGGCCAAGGAGGCCGGCCGGGCGCAAGCGCCACCGCCAGTACGCGCCTGCAGCAGTGGGGACCCCGCTGGGCGTTGCAACCCGTCGGGATG
TTGGGCGCTGCGTTTGTTCCCACCGACAGCGGTCAGCGAAATCCATCCTGCGGGTGAAAGCCGGTGTCCAGGCTAACCGGGATACAGGAGAAGGAGGGTAGCGAAGA
TGCTCAGTGCGTGGACAGTGCGTAACTTTGGTAATCGAGGCCGCACTGACGTGGTCCTGCCTCAGGCCACTCTCAGCCCCATCCTATCCCTGCTCGGGCCCGGTCCC
CCTCAGCTTGTCTTCATTTTTCGTCTCTATGAGTCATACTGAGAGCGACCTTCAGAAAGTGCGGTGGGTGGCGGTGGGTGGGTGGGTGGGGGGTTCAGGGGCGGGGA
ATACTGGCTGGTAGCCCCGCAGGATCAGATCACCAGGTTGTCCAGTGCACCCTGCGGGGCAGTGGCGAGCCTGGAGAGCCGCCTCTGCCTGGGAGAATTAACGACGT
CACAGCCAGAGACGAAAGAACTACAGGTCGCGGGGACAGGCTGGCTCCCAGGGTAGAGCTTGCAAAAGGGACGAGCGTGGGACAGACTGAGGTGGGACTGTTGTAGT
TCAGGGGCCGGCCGATCCGGCTCCGTAGGCTGCCCCTATGTCCCTCCCTCGCCCCATCCAGGCCAGCACAGCACGAGTAACGCCAGCCTAGAGCCCTCCCGCCGCGT
CCTCCCACTCCCGGCCCCACGTCTCGAAGCCACAGCCCAGCCCCACGCTTTGCCCGGCTGCACCCCGAGGTGCCCGCGTGCAGGGCGCGGGCCGGCAGGGCGCGCTT
GCCGACGCTCTTTTGTCTGCCAGTACCCAAAAAGCGGGGGGACGCGTGGCATCAGAATGTGGCAGATTGCGGAATATAGAAAAAAGAGGGTTACAGTTTTCTATGGG
AATGAGAGGTGCCAGGACTTGAAAGAAAATAAAACCACATACTCCCTCTGCTATATTAACTGAGAGTCCTTAAAACAAATAACTTCACGTTTA

### *PE Sox1(+35)TFBS* in *Gria1* promoter

GCCATCATTCTGTGCACACATCTCTCTTGGGTATCTCCGAGCCAGCTGCAGTCCAGCAGGGTCCACAGAGGCCCAGCTTCTCCTGGACACAAAC
AATCTGAGTTATAAGTGAAGGCAGCCTGCATCCTCCGACTGGAATCAGCGCAAAATGACTCATGTAATTGCCCTGTGTGCGTACCCTGAATCTT
CTTGTTCACCCACCCACACAACTCTGCTGTTATAGATTCCTGCACCCTGCAAGAAAAAAAAAGGGGAGGGGGGAGGACAAGTTCAAATACCTAT
GGTTGCAAAAAGGCCACCGACTAGTCCAGGAGCGTCGTGAGTTATGAGGTAGGAGGGAGCCGCTTGCCATTAATCTGGGAACAAACAGCTAACC
CCGGTGACTGGTATTTTTCCTTTTCTTTTCTCACTTTTCTCAGTGTGGGGAAGGCGATTGTGAGGCGCTCTGCTGGAATTTGGCAGCGCGGAGG
CTTGGAGAGCAGCCCCATGCTGGCTCCTATTCAGCCGGCCAGTTTTCCTCGAGCTTTGGAAGTTTCACTCAGCCGTGCACTCAATGGCTTCACA
AAGCTGATTACAAGCTTCAGCGCATTCCTGAAGGAGCCAAAAGCGACGCAGGTGCAAACGAGCCGAGGGAGCCCCTTATCCCGGTGACAGAATG
GGACAAGCTGGGAAAGGCTTGGACCACACAAATCCAAGGCTCACCAGGCAGCAGAGAGCCTGCCTTGGGAACCGGGGGTCATTATCCGCCCTAT
TCAGCGGGACCGGGGACCCTGGGGCCAAGGAGGCCGGCCGGGCCCAGGAGCGTCGTGAGTTATGAGACGCGTGCCTCTCCTAAAGATGGAAGGG
GCTGCTAGCATCCAGGTCCAAGCACAGGCCAGTCACACCGAAGCTACTGTTGTATGCCTTTCTCACAGTCTTTCCTTTAAATTTCTGGAGAGGG
ATTTCTAGGGTCTCCCCCTTGGGAATTAGTTGTAGGAATAATTGGGCCAGTGGAGTGTAGAAGATATATCCAGCGCAACCCGTGCGTGCACTAC
TAGAGAAAGTGACCTAGATCAAGCAGCTGGTGAATCCAGGGCTATTGTGG

### *PE Sox1(+35)TFBS+CGI* in Gria1 promoter

GCCATCATTCTGTGCACACATCTCTCTTGGGTATCTCCGAGCCAGCTGCAGTCCAGCAGGGTCCACAGAGGCCCAGCTTCTCCTGGACACAAAC
AATCTGAGTTATAAGTGAAGGCAGCCTGCATCCTCCGACTGGAATCAGCGCAAAATGACTCATGTAATTGCCCTGTGTGCGTACCCTGAATCTT
CTTGTTCACCCACCCACACAACTCTGCTGTTATAGATTCCTGCACCCTGCAAGAAAAAAAAAGGGGAGGGGGGAGGACAAGTTCAAATACCTAT
GGTTGCAAAAAGGCCACCGACTAGTCCAGGAGCGTCGTGAGTTATGAGGTGTAGGAGGGAGCCGCTTGCCATTAATCTGGGAACAAACAGCTAA
CCCCGGTGACTGGTATTTTTCCTTTTCTTTTCTCACTTTTCTCAGTGTGGGGAAGGCGATTGTGAGGCGCTCTGCTGGAATTTGGCAGCGCGGA
GGCTTGGAGAGCAGCCCCATGCTGGCTCCTATTCAGCCGGCCAGTTTTCCTCGAGCTTTGGAAGTTTCACTCAGCCGTGCACTCAATGGCTTCA
CAAAGCTGATTACAAGCTTCAGCGCATTCCTGAAGGAGCCAAAAGCGACGCAGGTGCAAACGAGCCGAGGGAGCCCCTTATCCCGGTGACAGAA
TGGGACAAGCTGGGAAAGGCTTGGACCACACAAATCCAAGGCTCACCAGGCAGCAGAGAGCCTGCCTTGGGAACCGGGGGTCATTATCCGCCCT
ATTCAGCGGGACCGGGGACCCTGGGGCCAAGGAGGCCGGCCGGGCGCAATCGATGTCATTATCCGCCCTATTCAGCGGGACCGGGGACCCTGGG
GCCAAGGAGGCCGGCCGGGCGCAAGCGCCACCGCCAGTACGCGCCTGCAGCAGTGGGGACCCCGCTGGGCGTTGCAACCCGTCGGGATGTTGGG
CGCTGCGTTTGTTCCCACCGACAGCGGTCAGCGAAATCCATCCTGCGGGTGAAAGCCGGTGTCCAGGCTAACCGGGATACAGGAGAAGGAGGGT
AGCGAAGATGCTCAGTGCGTGGACAGTGCGTAACTTTGGTAATCGAGGCCGCACTGACGTGGTCCTGCCTCAGGCCACTCTCAGCCCCATCCTA
TCCCTGCTCGGGCCCGGTCCCCCTCAGCTTGTCTTCATTTTTCGTCTCTATGAGTCATACTGAGAGCGACCTTCAGAAAGTGCGGTGGGTGGCG
GTGGGTGGGTGGGTGGGGGGTTCAGGGGCGGGGAATACTGGCTGGTAGCCCCGCAGGATCAGATCACCAGGTTGTCCAGTGCACCCTGCGGGGC
AGTGGCGAGCCTGGAGAGCCGCCTCTGCCTGGGAGAATTAACGACGTCACAGCCAGAGACGAAAGAACTACAGGTCGCGGGGACAGGCTGGCTC
CCAGGGTAGAGCTTGCAAAAGGGACGAGCGTGGGACAGACTGAGGTGGGACTGTTGTAGTTCAGGGGCCGGCCGATCCGGCTCCGTAGGCTGCC
CCTATGTCCCTCCCTCGCCCCATCCAGGCCAGCACAGCACGAGTAACGCCAGCCTAGAGCCCTCCCGCCGCGTCCTCCCACTCCCGGCCCCACG
TCTCGAAGCCACAGCCCAGCCCCACGCTTTGCCCGGCTGCACCCCGAGGTGCCCGCGTGCAGGGCGCGGGCCGGCAGGGCGCGCTTGCCGACGC
TCTTTTGTCTCCAGGAGCGTCGTGAGTTATGAGACGCGTGCCTCTCCTAAAGATGGAAGGGGCTGCTAGCATCCAGGTCCAAGCACAGGCCAGT
CACACCGAAGCTACTGTTGTATGCCTTTCTCACAGTCTTTCCTTTAAATTTCTGGAGAGGGATTTCTAGGGTCTCCCCCTTGGGAATTAGTTGT
AGGAATAATTGGGCCAGTGGAGTGTAGAAGATATATCCAGCGCAACCCGTGCGTGCACTACTAGAGAAAGTGACCTAGATCAAGCAGCTGGTGA
ATCCAGGGCTATTGTGG
